# Diet unmasks genetic variants that regulate lifespan in outbred *Drosophila*

**DOI:** 10.1101/2020.10.19.346312

**Authors:** Luisa F. Pallares, Amanda J. Lea, Clair Han, Elena V. Filippova, Peter Andolfatto, Julien F. Ayroles

## Abstract

Evolutionary theory suggests that lifespan-associated alleles should be purged from the gene pool, and yet decades of GWAS and model organism studies have shown they persist. Here, we address one potential explanation, the idea that the alleles that regulate lifespan do so only in certain contexts. We exposed thousands of outbred *Drosophil*a to a standard and a high sugar diet. We then sequenced over 10,000 individuals and track genome-wide allele frequency changes over time, as these populations aged. We mapped thousands of lifespan-altering alleles, some associated with early vs late life tradeoffs, late-onset effects, and genotype-by-environment interactions. We find that lifespan-reducing alleles are most likely to be recently derived, have stronger effects on a high-sugar diet, consistent with the hypothesis that historically neutral or beneficial alleles can become detrimental in novel conditions. We also show that the gene midway, a regulator of lipid storage and ortholog of the lifespan-associated gene DGAT1 in mice, also regulates lifespan in *Drosophila*. Our results provide insight into the highly polygenic and context-dependent genetic architecture of lifespan, as well as the evolutionary processes that shape this key trait.

## Introduction

Lifespan, a major component of fitness and a key life history trait, has a genetic basis: it is modestly heritable in humans and other organisms (h^2^~10%) (*1*) and dozens of lifespan-reducing alleles have now been identified (*2*, *3*). However, the fact that genetic variation for lifespan exists at all presents an evolutionary puzzle, as it is expected that natural selection will purge fitness-reducing alleles from the gene pool. Evolutionary theory provides several potential, non-mutually exclusive explanations for this conundrum. Lifespan-reducing alleles may persist because: (i) they are only deleterious in late-life, when selection is relatively weak (the mutation accumulation theory (*4*)), (ii) they provide benefits early in life that outweigh their late-life costs (the antagonistic pleiotropy theory (*5*)), and (iii) their effects vary across environments (genotype-by-environment, GxE) making them difficult to purge through purifying selection.

Notably, a special class of GxE interactions, driven by evolutionarily recent changes in human diet and lifestyle (*6–8*), are thought to be particularly important for human disease. Specifically, it has been proposed that many chronic, noncommunicable diseases are caused by alleles that evolved under stabilizing or positive selection throughout human history, but are now “mismatched” to obesogenic diets and other aspects of modern life (*6*–*9*). While this explanation is compelling, empirical data is limited due to the difficulty of identifying GxE interactions at genome-wide scale with high power (*10*). As a result, the degree to which exposure to evolutionarily novel environments alters the relationship between genetic variation and fitness-related traits remains unclear.

To address this question, we leveraged the tractable experimental and genomic tools of *Drosophila melanogaster* to map loci that affect lifespan in two environments. Specifically, we exposed an outbred population of flies to two diets: a standard laboratory diet and a high sugar diet containing more sucrose than flies would encounter in nature and that is known to cause obesity, diabetes, and reduced lifespan in this species (*11*, *12*). Drawing inspiration from a recent study of human longevity (*13*), we tracked genome-wide allele frequency changes in age-matched flies across their entire adult life. Using this high-powered experimental approach (Figure 1A-B), we were able to identify thousands of lifespan-reducing alleles that decrease in frequency as individuals grow older, as well as to classify them into: (i) late-onset alleles that only decrease at late ages (mutation accumulation theory), (ii) alleles with tradeoffs between early and late life that first increase and then decrease (antagonistic pleiotropy theory), and (iii) GxE alleles that have substantially stronger effects on lifespan on one diet, potentially due to risk alleles being exposed by the novel diet. Together, our study provides insight into the genetic architecture and environmental sensitivity of a major life history trait, and experimentally evaluates evidence for long-standing theories for why fitness-reducing alleles abound in nature.

**Fig. 1.**
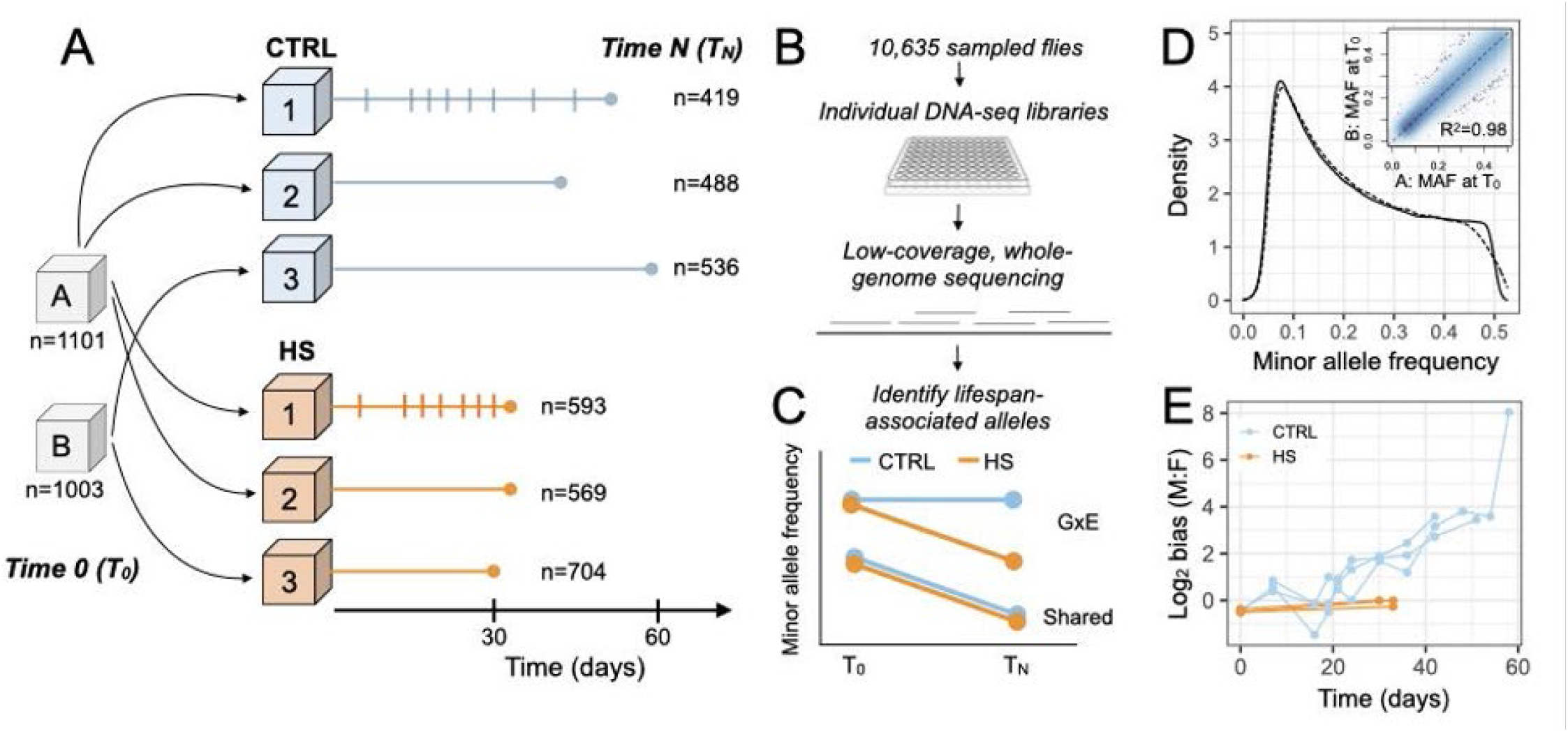
Experimental design to detect GxE interactions modulating lifespan. (**A**) *D. melanogaster* flies caught in Princeton, NJ were used to generate a synthetic outbred population that was kept under laboratory conditions for over a year, and split in two replicate cages prior to the beginning of the experiment (A and B). ~1000 flies were collected from A and B at the start of the experiment (*T_0_*, 2±1 days old) and the rest were distributed into 6 replicate cages of ~10,000 flies each (3 cages = standard lab diet (CTRL), blue; 3 cages = high sugar diet, orange). ~500 flies were sampled every 3-7 days from a given cage and a last sample was taken when only ~500 flies were left (*T_N_*) (Table S1-2); sampling schedule are noted by vertical dashed lines for the CTRL1 and HS1 cages. Identical schedules were followed for all other cages within a treatment group. To prevent pupae from the new generation from eclosing inside the experimental cages, food containers were replaced every three days. (**B**) Individually barcoded DNA-seq libraries were prepared from 10,635 individual flies sampled from *T_0_*, *T_N_*, and the intermediate time points. Each library was sequenced at ~1x depth to estimate allele frequencies and test for frequency changes across time (Fig. S1). **(C)** Expected patterns of frequency change are shown for alleles that reduce lifespan in both diets (shared) or more so on the HS diet (GxE). (**D**) The allelic composition of cages A (solid line) and B (dashed line) is very similar at *T_0_* (n=291,319 SNPs). Inset shows the per-site correlation between the minor allele frequency (MAF) estimated for cage A versus B at *T_0_*. (**E**) Log_2_ ratio of males to females at different timepoints during the experiment. The number of flies sexed at each time point is provided in Table S4.

### Sex and genotype have environment-specific effects on lifespan

To identify loci associated with lifespan variation and evaluate their context-dependence, we exposed large, replicate populations of age-matched outbred adult flies to a standard laboratory diet (hereafter “control” or “CTRL”) and a high sugar diet (“HS”) for one generation (n=3 replicates of ~10,000 flies per diet; Figure 1A). To prevent overlapping generations, food containers (where flies also lay eggs) were exchanged every three days. We drew a random sample of ~2000 flies at the beginning of the experiment (*T_0_*), and continued to sample ~500 flies from each population at regular intervals. When only the ~500 longest-lived flies remained in a given replicate cage, we collected a final sample (*T_N_*) (Table S1,2). In total, 10,637 flies were genotyped using individually barcoded low-coverage genome sequencing (Fig. S1), and used to estimate age-specific genome-wide allele frequencies on each diet.

While all replicates for the two diets started from a common pool of standing genetic variation at the beginning of the experiment (Figure 1D; Table S3), we observed a consistent, ~1.6 fold reduction in lifespan for flies on the HS diet, as expected (*12*). We also observed substantial and unexpected interactions between diet and sex: while the sex ratio remained roughly 1:1 as flies aged on the HS diet, males far outlived females on the CTRL diet resulting in a sex ratio of ~100:1 by the end of the experiment (Figure 1E; Table S4). We replicated this observation in independent experiments where the lifespan of individual flies was quantified, suggesting that it is a repeatable characteristic of the fly population used here (Cox proportional hazards: p(sex-by-diet) = 0.026) (Fig. S2, Table S5). While others have also observed that sex-specific lifespans in flies are sometimes environmentally-dependent (*14*, *15*), future work is necessary to uncover the proximate mechanisms at play here.

To detect longevity-associated alleles, we estimated allele frequencies at 268,159 common SNPs (MAF>0.05) and tested for alleles that exhibited a significantly lower frequency at the end of the experiment (*T_N_*; n=1443 and 1866 sequenced flies for CTRL and HS, respectively) compared to the beginning of the experiment (*T_0_*; n=2104 sequenced flies). Such decreases in frequency indicate that individuals carrying a given allele die at younger ages relative to individuals carrying the alternative allele (Figure 1C). We identified 2246 genetic variants that fit this pattern, distributed among 1919 genes (permutation-derived FDR 10%; Figure 2, Table S6, Fig. S3,4). The average absolute decrease in allele frequency between *T_N_* and *T_0_* for these lifespan-associated SNPs was 0.08, with most changes falling between 0.05-0.12 (Fig. S5). The majority of the lifespan-associated SNPs (68.6%) have significant and positively correlated effects on the two diets, suggesting similar or “shared” effects are common (Figure 2C, Fig S4). However, we found that 31.4% of lifespan-associated SNPs (n=704) exhibited evidence for substantial GxE interactions (i.e., stronger effects on one diet relative to the other, defined as permutation derived FDR<10% in one environment and p>0.05 in the other environment, see also Fig. S6). Strikingly, of these 704 SNPs with the largest GxE effects on lifespan, 99.6% (701) had larger effects on the HS diet, indicating that their effects are magnified or unmasked under dietary stress (Figure 2). These results suggest that a substantial amount of genetic variation that appears to have little effect on phenotypic variation under one set of conditions might indeed play a fundamental role in new or stressful environments. We also note that SNPs identified as having “shared” effects in the two environments more often exhibit stronger effects on the HS diet (76% of the time, p-value = 1.4e-98). These results, and our simulations (Fig S6), suggest that GxE interaction is a common feature among SNPs affecting lifespan.

**Fig. 2.**
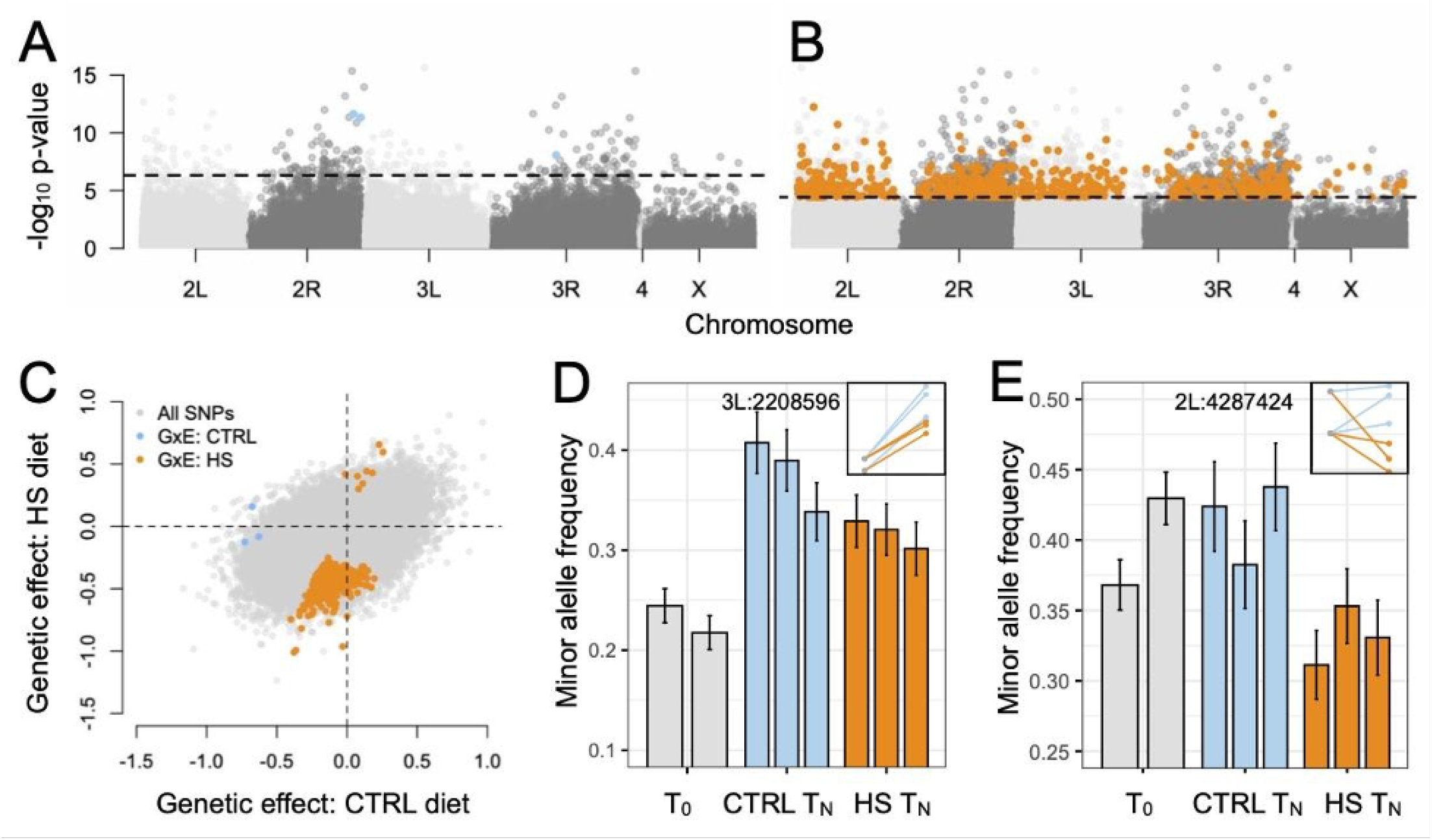
GxE interactions determine lifespan. (**A**, **B**) Manhattan plots highlighting significant lifespan-associated SNPs with the strongest GxE effects. Plots show the −log_10_ p-value for tests for allele frequency differences between *T_N_* and *T_0_* on a (A) CTRL and (B) HS diet; colored points passed our significance filters for GxE effects (see methods). The p-value threshold corresponding to a 10% empirically determined FDR is noted with a dashed line for each environment. (**C**) Comparison of model-estimated effect sizes for a genetic effect on lifespan on CTRL versus HS diets (positive values indicate the alternate allele increases in frequency at *T_N_* versus *T_0_*). Only SNPs with significant evidence for GxE effects are colored. (**D, E**) Allele frequency changes across replicates for (D) an example SNP (3L:2208596) associated with lifespan in both dietary conditions and (E) an example SNP (2L:4287424) with larger effects on lifespan on the HS diet. The estimated minor allele frequency is shown for each replicate cage, with bars representing the standard error. The two T_0_ bars correspond to cage A and B. The inset shows the mean minor allele frequencies at *T_N_* and *T_0_*, for each replicate CTRL (blue) and HS (orange) cage, using the same x and y axes as in Figure 1B.

### Biological and functional insight into the genetic basis of lifespan

To understand the biology of loci that contribute to lifespan we first asked whether lifespan-associated SNPs were enriched in particular genomic features or molecular processes. We found that our longevity-associated SNPs are not significantly enriched for any particular molecular pathway nor for “canonical” longevity genes (Table S7). These results support a highly polygenic model in which genetic variation segregating in wild-derived populations of *D. melanogaster* does not localize to the canonical biological pathways associated with aging and lifespan (*16*, *17*), as was also observed by (*14*, *18*, *19*). Notably, we did find that lifespan-associated SNPs are strongly enriched in genes identified in previous studies of *D. melanogaster* longevity (Figure 3A, Table S8, S9).

**Fig. 3.**
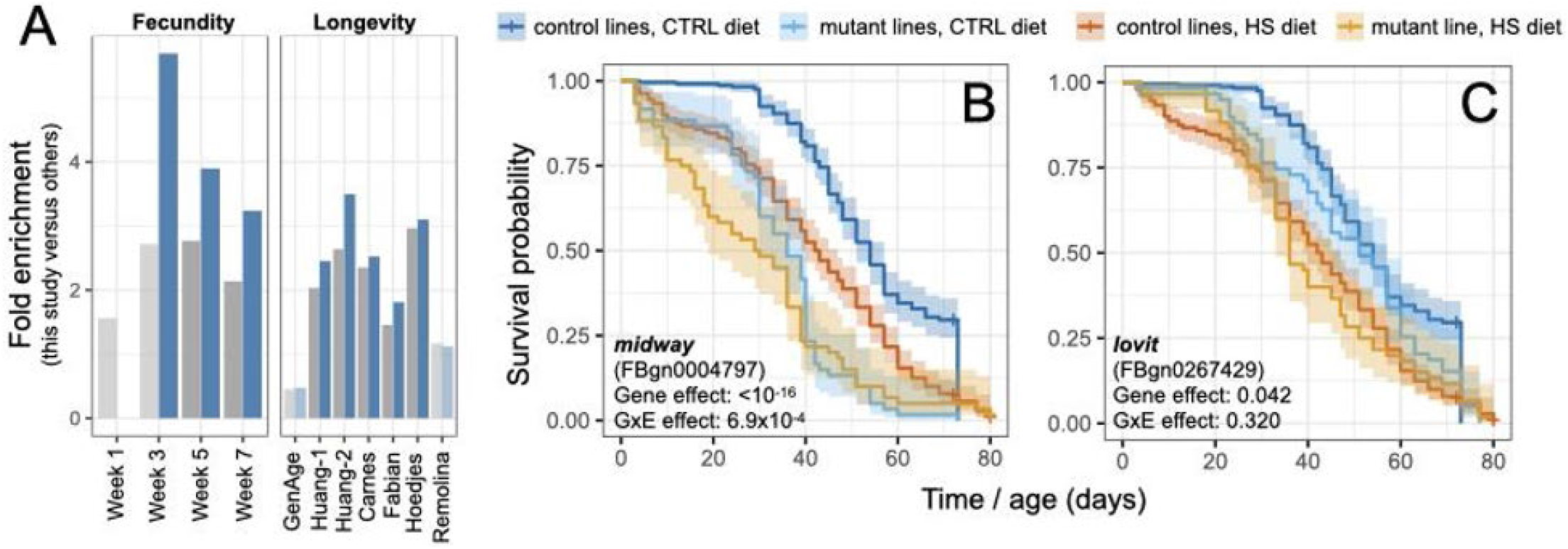
Properties of lifespan-associated genes. (**A**) Genes in or near (<1kb) SNPs with shared (grey) or substantial GxE (blue) effects on lifespan in this experiment overlap with lifespan and fecundity genes identified in previous studies (identified by first author’s last name). The degree of overlap is represented as fold enrichment from a Fisher’s exact test, and light bars indicate non-significant overlap. Studies represent several types: GWAS for fecundity measured during weeks 1-7 in inbred lines (*25*); selection for extended lifespan in outbred flies (*18*, *19*, *26*, *27*); analyses of standing variation associated with lifespan in inbred lines (*14*); and “canonical” longevity genes from the GenAge database (*28*). Light bars indicate non-significant overlap. (**B, C**) Kaplan-Meier survival curves for two candidate genes, with p-values from a Cox proportional-hazards model testing for an effect of the gene on survival as well as a GxE effect. Survival curves for the control lines include data from four wild-type control lines (DGRP 439, DGRP 181, Canton-S, and yw).

Many of the lifespan-associated genes we identified perform essential functions but are not known to affect lifespan in flies. For example, *midway*, involved in fat metabolism and oogenesis (*20*), and *lovit*, involved in neurophysiology (*21*), both contain lifespan-associated SNPs. Using loss-of-function mutant lines, we validated their effects on lifespan (*lovit*: p-value = 0.042; *midway*: p-value < 10^−16^, Figure 3B-C, Table S10). The validation of *midway*, a diacylglycerol acyltransferase involved in triglyceride metabolism, is a notable finding since its mammalian ortholog, *DGAT1*, has been shown to regulate lifespan in mice (*22*). In addition, *midway* has a particularly strong GxE effect in the outbred population (change in MAF with age on HS = 12.5%, q-value = 0.04; on CTRL = 3.8%, q-value = 0.7) as well as in the loss-of-function lines (p-value for GxE interaction = 7×10^−4^, Figure 3B-C, Table S10), indicating that its effect on lifespan is environment-dependent. These results contribute to the increasing body of literature linking lipid metabolism to the regulation of aging and lifespan (*23*).

### Testing evolutionary theories of aging and longevity

Fitness-reducing alleles are thought to be largely governed by mutation-selection balance, in which mutation continuously generates deleterious alleles and purifying selection eliminates them (*24*). However, we find that first, the minor allele reduces lifespan only in about half of cases (58%), and second, that these risk alleles are by no means rare in the population (mean frequency +/- SD at T_0_ = 0.35 +/- 0.1; Table S6), suggesting that other evolutionary forces maintain them at moderate frequencies. Our results indicate that GxE interactions are one key factor. We next asked if two additional forces, antagonistic pleiotropy and mutation accumulation, may also be important contributors to this feature of the data. Specifically, we estimated allele frequencies at several time points between *T_0_* and *T_N_* to determine the trajectory of lifespan-reducing alleles (Figure 1A, Table S1, S2). We then asked whether these alleles exhibited (i) a U-shaped pattern indicative of trade-offs and differential fitness effects at young versus old ages, as predicted by antagonistic pleiotropy theory (*5*); (ii) evidence for fitness-effects only at old ages, as predicted by mutation accumulation theory (*4*); or (iii) an evolutionary “null” model of constant fitness-effects at all ages (Figure 4A).

**Fig. 4.**
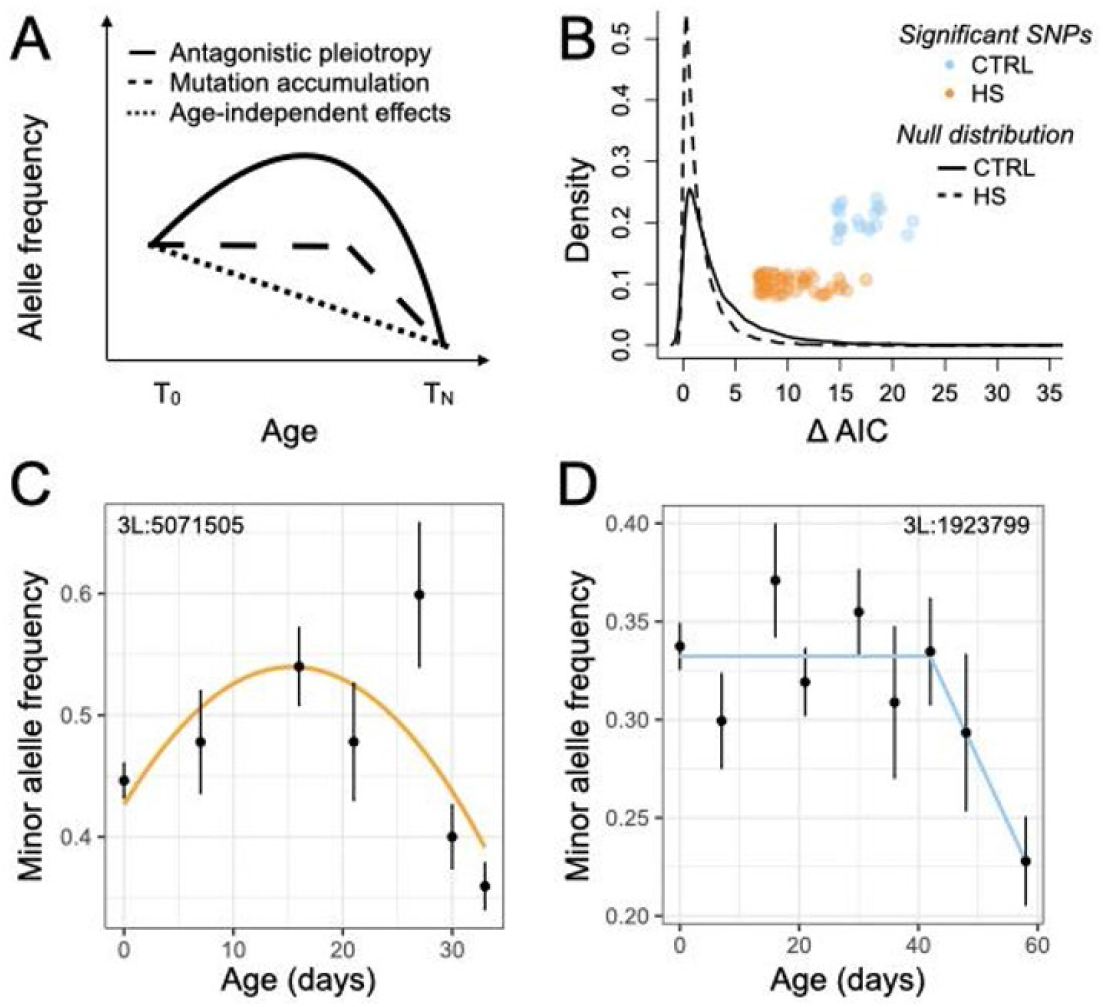
Insights into evolutionary theories of aging. **(A)** Allele frequency trajectories across time according to the antagonistic pleiotropy and mutation accumulation theories, and a constant trajectory not expected under evolutionary models. We asked whether each lifespan-associated SNP could be confidently assigned to one of these trajectories. (**B**) ΔAIC between the best and second-best model for each tested SNP. HS and CTRL cages were analyzed separately due to the different age distributions within each treatment (See Figure 1A). SNPs with ΔAIC values >97.5% of the null distribution are confidently assigned to a given trajectory and their ΔAIC values are plotted as individual points. Examples of a **(C)** quadratic trajectory SNP in HS suggesting antagonistic pleiotropy and **(D)** a breakpoint trajectory SNP in the standard lab environment (CTRL) suggesting mutation accumulation dynamics. Points represent the mean alternate allele frequency for a given age estimated across all cages, while bars represent the standard error of the estimate.

Of the 2246 SNPs with significant effects on lifespan, we confidently assigned 75 to one of the three trajectories described above (ΔAIC between the best and second-best trajectory > 97.5% of permutations). 29 (42%) of these SNPs exhibit an antagonistic pleiotropy pattern, and 38 (48%) exhibit a pattern consistent with mutation accumulation theory (Figure 4B-D; Table S11). In further support of antagonistic pleiotropy theory, we also find that genes near lifespan-associated SNPs (not just the 75 with assigned trajectories) significantly overlap with genes identified in a previous study of age-specific fecundity in flies (Figure 3A; Table S9; (*25*)). Interestingly, the pattern is most pronounced for the 704 SNPs we identified with the strongest GxE effects (Figure 3A, Table S9). This overlap further indicates that many longevity-decreasing alleles are maintained because they provide other benefits, for example to fertility in early adulthood, that outweigh their late life costs.

### The evolution of alleles regulating lifespan

The finding that GxE interactions are common with respect to diet in our experiment has important implications for human health. In particular, it is thought that rapid shifts in human diet and lifestyle following the Industrial Revolution have caused previously adaptive or neutral alleles to become maladaptive (or “mismatched”), such that they are currently associated with diseases that impact lifespan (Figure 5A; (*6*, *7*, *10*)). The high-sugar (HS) environment provided in our experiment is a particularly extreme case of such a change, and potentially relevant to relatively recent dietary changes in some human populations. We have shown that exposing flies to such high sugar concentrations reveals a substantial amount of genetic variation that remains hidden/cryptic in the CTRL diet, as has been predicted repeatedly (6) but rarely tested experimentally. That said, it should also be noted that, from an evolutionary perspective, our standard lab media (CTRL) also reflects a substantial change in diet from that experienced by the wild-caught flies in our experiment (collected in Princeton, NJ) and the ancestral populations from which they ultimately originated (sub-Saharan Africa). The prevalence of GxE interactions among the SNPs we have associated with lifespan allows us to test predictions of the “evolutionary mismatch” hypothesis for the first time at genome-wide scale. Specifically, we asked whether alleles with lifespan-reducing effects in the Princeton population 1) are more likely to be recently derived and 2) exhibit signatures of positive selection in this population and/or the ancestral populations from which it originated.

**Fig. 5.**
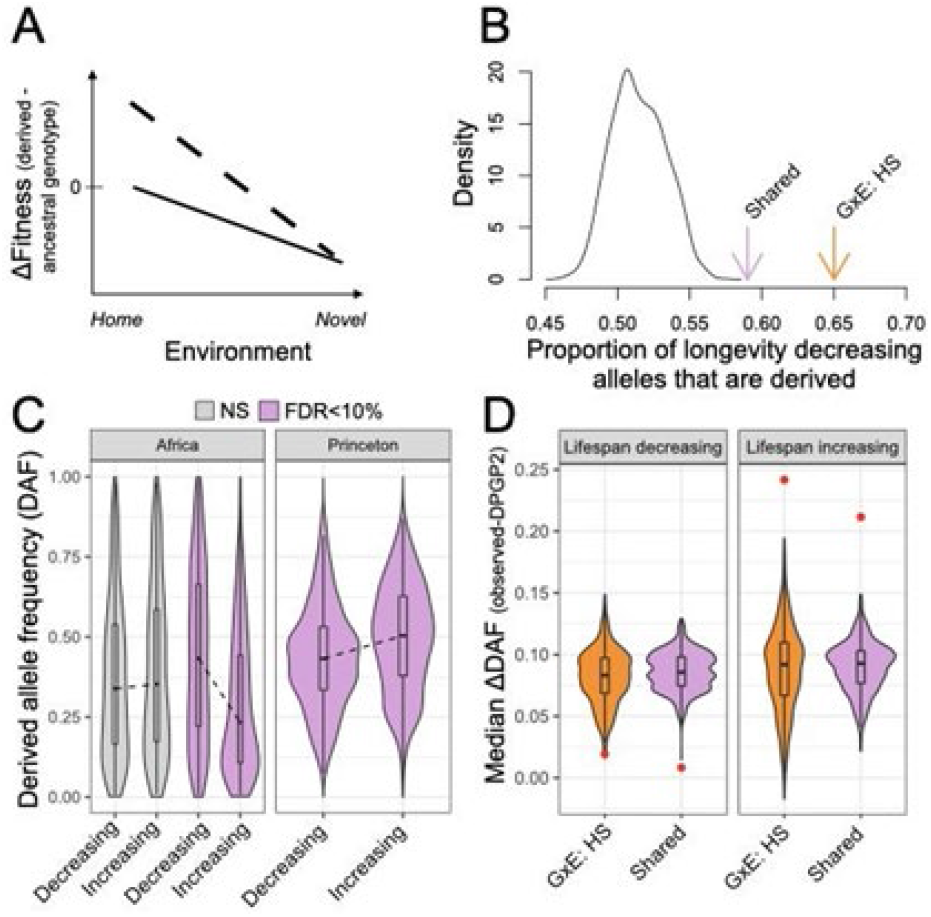
Insights into the evolution of alleles that regulate lifespan. **(A)** Potential predictions from mismatch theory: alleles that evolved more recently in the focal population (derived alleles) are neutral (black line) or advantageous (dashed line) in the ancestral environment they evolved in; however, they become deleterious in a “novel” environment. **(B)** The derived allele is more likely than the ancestral allele to reduce lifespan relative to chance expectations. For lifespan-associated SNPs shared between environments (Shared) or with substantially stronger effects on the HS diet (GxE:HS), the proportion of SNPs for which the derived allele is the lifespan-reducing allele is noted with an arrow. Null expectations were derived by performing the same calculations on effect sizes from individual CTRL and HS cages across 1000 randomly drawn pools of 1000 frequency matched non-significant SNPs. **(C)** The derived allele frequency (DAF) distribution for sites where the derived allele significantly increases or decreases lifespan in at least one environment (FDR<10%). For comparison, the DAF for a set of non-significant sites (NS) frequency-matched to lifespan-increasing and decreasing alleles in Princeton is also shown. The DAF distribution is plotted for our experiment at T0 (“Princeton”) as well as for the DPGP2 African dataset (“Africa”) for the same set of sites. Lines highlight the differences in median values between each pair of distributions. DAF for lifespan-increasing and decreasing alleles in the Princeton as well as in the African populations are significantly different, but with opposite directions (Wilcoxon test: Princeton p-value = 5e-8, Africa p-value = 4e-16). The distributions of NS sites are not significantly different. **(D)** DAF difference in our experiment (at T0) versus the DPGP2 for lifespan increasing and decreasing alleles. Red dots indicate the median DAF difference observed in real data for a given set of alleles. Colored distributions represent the median DAF differences for 1000 datasets of non-significant sites sampled to match the frequency distribution of the allele set.

As predicted, we find that lifespan-reducing alleles are more likely to be derived than ancestral, and this bias is the largest for SNPs with the strongest GxE effects (65%, Figure 5B). We next asked if lifespan-associated alleles exhibit evidence for selection, and of what form. Notably, we find that lifespan-decreasing alleles are at significantly lower frequency than lifespan-increasing alleles in the Princeton population (p-value = 8e-8, Wilcoxon test; Figure 5C), consistent with their predicted effects on fitness. To further elucidate the nature of selection pressures on lifespan-associated alleles, we determined the frequencies of these alleles and frequency-matched non-significant SNPs in putatively ancestral African populations (Figure 5C). Remarkably, these comparisons reveal that, in contrast with what is observed in the Princeton population, in Africa lifespan-decreasing alleles are at significantly higher frequency than lifespan-increasing alleles (p-value = 4e-16, Wilcoxon test). Moreover, lifespan-decreasing alleles are at significantly higher frequencies than non-significant SNPs in African populations (p-value = 2e-5, Wilcoxon test), consistent with positive selection, either direct or via linked-selection, promoting lifespan-decreasing alleles in these populations (the frequencies of non-significant SNPs for lifespan-increasing and lifespan-decreasing alleles do not differ, p-value = 0.24, Wilcoxon test, Figure 5C). When Princeton and African allele frequencies are directly compared, we observe that while lifespan-decreasing alleles do not differ in frequency between Princeton and African populations, lifespan-increasing alleles show a notable increase in frequency in the Princeton population (Figure 5D). The differences between Princeton and African populations suggest that spatially and/or temporally heterogeneous selection pressures have impacted the frequencies of lifespan-associated SNPs. The evidence for positive selection in the evolutionary history of alleles that decrease lifespan in the Princeton population supports the evolutionary mismatch hypothesis for the presence of high frequency alleles that are detrimental to fitness when exposed to novel environments.

### Implications for understanding the genetic basis of lifespan variation

Long-standing population genetic and evolutionary theories have proposed several forces at play in the maintenance of genetic variation for fitness-related traits (*4*, *6*–*8*). Experimental tests of the predictions of these theories have been hampered by the difficulty of mapping fitness-related genetic effects. For example, a recent human study using a similar approach to ours but a 10-fold larger sample size found only two lifespan-associated regions near the *APOE* and *CHRNA3* genes (*13*). Instead, we identified thousands of lifespan-associated loci, most of which have larger effects on the HS diet, uncovering a highly polygenic and context-dependent architecture. We estimate that in the absence of environmental heterogeneity, both studies have similar statistical power (Fig. S7); the fact that we find orders of magnitude more lifespan-associated SNPs here highlights the utility of well-controlled experimental designs in model organisms for the study of complex traits. Because our high-powered design allowed us to identify many lifespan-reducing alleles, we could evaluate the generality of important theories for why alleles that shorten lifespan persist in nature and how they evolve. Specifically, we identified a key role for GxE interactions, as well as mutation accumulation and antagonistic pleiotropy as forces maintaining genetic variation for lifespan. We also provide experimental insight into how interactions between derived genetic variation and novel environmental conditions may shorten lifespan.

## ACKNOWLEDGMENTS

We thank Andrew M. Taverner for providing the multi-species alignments, Erika Gadja for help collecting flies from the wild, and Michael Fernandez for help maintaining the experimental cages. We also thank members of the Ayroles and Andolfatto labs for their general support and helpful scientific discussions. AG Clark, M. Przeworski, G. Sella and A. Harpak provided helpful comments on the analyses.

## Author contributions

LFP: conceptualization, formal analysis, investigation, writing-original draft, writing-review and editing; AJL: conceptualization, formal analysis, investigation, writing-review and editing, visualization; CH: conceptualization, formal analysis, investigation, writing-review and editing; EVF: investigation; PA: conceptualization, formal analysis, writing-review and editing, funding acquisition; JFA: conceptualization, formal analysis, writing-review and editing, funding acquisition

## Competing interests

Authors declare no competing interests.

### Data and materials availability

the data and code for analysis will be made available after publication.

## SUPPLEMENTARY MATERIALS

- Materials and Methods. References cited only in this section (*29*–*59*)
- Figure S1. Mean read coverage for DNA libraries.
- Figure S2. Sex and diet interact to affect survival.
- Figure S3. QQ-plots comparing empirical and observed distribution of p-values for the genetic effect on longevity.
- Figure S4. Comparison of the magnitude of the genetic effect on longevity across conditions.
- Figure S5. Allele frequency changes between T0 and TN.
- Figure S6. Simulations to understand our definitions of “shared” and “GxE” SNPs.
- Figure S7. Power to detect a genetic effect on longevity using different study designs.
- Figure S8. Sex determined from sequence data.
- Figure S9. Power comparison between a Cochran-Mantel-Haenszel (CMH) and beta-binomial approach.
- Table S1. Sampling schedule by time point and cage.
- Table S2. Number of flies sequenced and analyzed per time point and cage
- Table S3. Fst and Pi calculations
- Table S4. Sex bias by time point and cage.
- Table S5. Results from a Cox proportional hazards mixed effects model, testing for condition and sex 5effects on lifespan in vials
- Table S6. Sites with significant (at a 10% FDR) genetic effects on at least one diet
- Table S7. Enrichment of SNPs with significant genetic effects in genomic features
- Table S8. Genes related to fecundity and longevity (from this study and others).
- Table S9. Enrichment of genes near SNPs with significant genetic effects on longevity in previously published data sets.
- Table S10. Results from a Cox proportional hazards mixed effects model, testing for sex, condition, and genotype effects on lifespan.
- Table S11. Sites for which an allele frequency trajectory could be confidently assigned.
- Table S12. List of SNPs for which ancestral state could be confidently assigned

## REFERENCES

1. J. G. Ruby, K. M. Wright, K. A. Rand, A. Kermany, K. Noto, D. Curtis, N. Varner, D. Garrigan, D. Slinkov, I. Dorfman, J. M. Granka, J. Byrnes, N. Myres, C. Ball, Estimates of the Heritability of Human Longevity Are Substantially Inflated due to Assortative Mating. Genetics. 210, 1109–1124 (2018).

2. M. Kuningas, S. P. Mooijaart, D. V. Heemst, B. J. Zwaan, P. E. Slagboom, R. G. J. Westendorp, Genes encoding longevity: from model organisms to humans. Aging Cell. 7, 270–280 (2008).

3. P. K. Joshi, K. Fischer, K. E. Schraut, H. Campbell, T. Esko, J. F. Wilson, Variants near CHRNA3/5 and APOE have age- and sex-related effects on human lifespan. Nature Communications. 7, 11174 (2016).

4. P. B. Medawar, An unsolved problem of biology (Published for the College by H.K. Lewis, London, 1952).

5. G. C. Williams, Pleiotropy, Natural Selection, and the Evolution of Senescence. Evolution. 11, 398–411 (1957).

6. G. Gibson, Decanalization and the origin of complex disease. Nature Reviews Genetics. 10, 134–140 (2009).

7. S. Corbett, A. Courtiol, V. Lummaa, J. Moorad, S. Stearns, The transition to modernity and chronic disease: mismatch and natural selection. Nature Reviews Genetics. 19, 419–430 (2018).

8. G. Gibson, I. Dworkin, Uncovering cryptic genetic variation. Nature Reviews Genetics. 5, 681–690 (2004).

9. A. Di Rienzo, R. R. Hudson, An evolutionary framework for common diseases: the ancestral-susceptibility model. Trends in Genetics. 21, 596–601 (2005).

10. L. Quintana-Murci, Understanding rare and common diseases in the context of human evolution. Genome Biology. 17, 225 (2016).

11. L. P. Musselman, J. L. Fink, K. Narzinski, P. V. Ramachandran, S. S. Hathiramani, R. L. Cagan, T. J. Baranski, A high-sugar diet produces obesity and insulin resistance in wild-type Drosophila. Disease Models & Mechanisms. 4, 842–849 (2011).

12. J. Na, L. P. Musselman, J. Pendse, T. J. Baranski, R. Bodmer, K. Ocorr, R. Cagan, A Drosophila Model of High Sugar Diet-Induced Cardiomyopathy. PLOS Genetics. 9, e1003175 (2013).

13. H. Mostafavi, T. Berisa, F. R. Day, J. R. B. Perry, M. Przeworski, J. K. Pickrell, Identifying genetic variants that affect viability in large cohorts. PLOS Biology. 15, e2002458 (2017).

14. W. Huang, T. Campbell, M. A. Carbone, W. E. Jones, D. Unselt, R. R. H. Anholt, T. F. C. Mackay, Context-dependent genetic architecture of Drosophila life span. PLoS Biol. 18, e3000645 (2020).

15. J. M. S. Burger, D. E. L. Promislow, Sex-Specific Effects of Interventions That Extend Fly Life Span. Sci. Aging Knowl. Environ. 2004, pe30 (2004).

16. M. D. W. Piper, L. Partridge, Drosophila as a model for ageing. Biochimica et Biophysica Acta (BBA) - Molecular Basis of Disease. 1864, 2707–2717 (2018).

17. C. López-Otín, M. A. Blasco, L. Partridge, M. Serrano, G. Kroemer, The Hallmarks of Aging. Cell. 153, 1194–1217 (2013).

18. K. M. Hoedjes, J. van den Heuvel, M. Kapun, L. Keller, T. Flatt, B. J. Zwaan, Distinct genomic signals of lifespan and life history evolution in response to postponed reproduction and larval diet in Drosophila. Evol Lett. 3, 598–609 (2019).

19. D. K. Fabian, K. Garschall, P. Klepsatel, G. Santos-Matos, É. Sucena, M. Kapun, B. Lemaitre, C. Schlötterer, R. Arking, T. Flatt, Evolution of longevity improves immunity in Drosophila. Evolution Letters. 2, 567–579 (2018).

20. M. Buszczak, X. Lu, W. A. Segraves, T. Y. Chang, L. Cooley, Mutations in the midway gene disrupt a Drosophila acyl coenzyme A: diacylglycerol acyltransferase. Genetics. 160, 1511–1518 (2002).

21. Y. Xu, T. Wang, LOVIT Is a Putative Vesicular Histamine Transporter Required in Drosophila for Vision. Cell Reports. 27, 1327–1333.e3 (2019).

22. R. S. Streeper, C. A. Grueter, N. Salomonis, S. Cases, M. C. Levin, S. K. Koliwad, P. Zhou, M. D. Hirschey, E. Verdin, R. V. Farese, Deficiency of the lipid synthesis enzyme, DGAT1, extends longevity in mice. Aging (Albany NY). 4, 13–27 (2012).

23. A. A. Johnson, A. Stolzing, The role of lipid metabolism in aging, lifespan regulation, and age-related disease. Aging Cell. 18, e13048 (2019).

24. J. F. Crow, M. Kimura, An Introduction to Population Genetics Theory (Blackburn Press, 2009).

25. M. F. Durham, M. M. Magwire, E. A. Stone, J. Leips, Genome-wide analysis in Drosophila reveals age-specific effects of SNPs on fitness traits. Nature Communications. 5, 4338 (2014).

26. M. U. Carnes, T. Campbell, W. Huang, D. G. Butler, M. A. Carbone, L. H. Duncan, S. V. Harbajan, E. M. King, K. R. Peterson, A. Weitzel, S. Zhou, T. F. C. Mackay, The Genomic Basis of Postponed Senescence in Drosophila melanogaster. PLOS ONE. 10, e0138569 (2015).

27. S. C. Remolina, P. L. Chang, J. Leips, S. V. Nuzhdin, K. A. Hughes, Genomic basis of aging and life-history evolution in Drosophila melanogaster. Evolution. 66, 3390–3403 (2012).

28. R. Tacutu, D. Thornton, E. Johnson, A. Budovsky, D. Barardo, T. Craig, E. Diana, G. Lehmann, D. Toren, J. Wang, V. E. Fraifeld, J. P. de Magalhães, Human Ageing Genomic Resources: new and updated databases. Nucleic Acids Res. 46, D1083–D1090 (2018).

29. T. M. Therneau, coxme: Mixed Effects Cox Models (2020; https://CRAN.R-project.org/package=coxme).

30. T. M. Therneau, P. M. Grambsch, Modeling Survival Data: Extending the Cox Model (Springer-Verlag, New York, 2000; https://www.springer.com/gp/book/9780387987842), Statistics for Biology and Health.

31. T. M. Therneau, T. L. (original S.->R port and R. maintainer until 2009), A. Elizabeth, C. Cynthia, survival: Survival Analysis (2020; https://CRAN.R-project.org/package=survival).

32. S. Picelli, A. K. Björklund, B. Reinius, S. Sagasser, G. Winberg, R. Sandberg, Tn5 transposase and tagmentation procedures for massively scaled sequencing projects. Genome Res. 24, 2033–2040 (2014).

33. M. Martin, Cutadapt removes adapter sequences from high-throughput sequencing reads. EMBnet.journal. 17, 10–12 (2011).

34. H. Li, R. Durbin, Fast and accurate short read alignment with Burrows-Wheeler transform. Bioinformatics. 25, 1754–1760 (2009).

35. G. A. Auwera, M. O. Carneiro, C. Hartl, R. Poplin, G. del Angel, A. Levy-Moonshine, T. Jordan, K. Shakir, D. Roazen, J. Thibault, From FastQ data to high-confidence variant calls: the genome analysis toolkit best practices pipeline. Curr. Protoc. Bioinformatics, 11–10.

36. S. Purcell, B. Neale, K. Todd-Brown, L. Thomas, M. A. R. Ferreira, D. Bender, J. Maller, P. Sklar, P. I. W. de Bakker, M. J. Daly, P. C. Sham, PLINK: a tool set for whole-genome association and population-based linkage analyses. Am. J. Hum. Genet. 81, 559–575 (2007).

37. B. S. Weir, C. C. Cockerham, Estimating F-Statistics for the Analysis of Population Structure. Evolution. 38, 1358–1370 (1984).

38. R. M. Durbin, D. Altshuler, R. M. Durbin, G. R. Abecasis, D. R. Bentley, A. Chakravarti, A. G. Clark, F. S. Collins, F. M. De La Vega, P. Donnelly, M. Egholm, P. Flicek, S. B. Gabriel, R. A. Gibbs, B. M. Knoppers, E. S. Lander, H. Lehrach, E. R. Mardis, G. A. McVean, D. A. Nickerson, L. Peltonen, A. J. Schafer, S. T. Sherry, J. Wang, R. K. Wilson, R. A. Gibbs, D. Deiros, M. Metzker, D. Muzny, J. Reid, D. Wheeler, J. Wang, J. Li, M. Jian, G. Li, R. Li, H. Liang, G. Tian, B. Wang, J. Wang, W. Wang, H. Yang, X. Zhang, H. Zheng, E. S. Lander, D. Altshuler, L. Ambrogio, T. Bloom, K. Cibulskis, T. J. Fennell, S. B. Gabriel, D. B. Jaffe, E. Shefler, C. L. Sougnez, D. R. Bentley, N. Gormley, S. Humphray, Z. Kingsbury, P. Kokko-Gonzales, J. Stone, K. J. McKernan, G. L. Costa, J. K. Ichikawa, C. C. Lee, R. Sudbrak, H. Lehrach, T. A. Borodina, A. Dahl, A. N. Davydov, P. Marquardt, F. Mertes, W. Nietfeld, P. Rosenstiel, S. Schreiber, A. V. Soldatov, B. Timmermann, M. Tolzmann, M. Egholm, J. Affourtit, D. Ashworth, S. Attiya, M. Bachorski, E. Buglione, A. Burke, A. Caprio, C. Celone, S. Clark, D. Conners, B. Desany, L. Gu, L. Guccione, K. Kao, A. Kebbel, J. Knowlton, M. Labrecque, L. McDade, C. Mealmaker, M. Minderman, A. Nawrocki, F. Niazi, K. Pareja, R. Ramenani, D. Riches, W. Song, C. Turcotte, S. Wang, E. R. Mardis, R. K. Wilson, D. Dooling, L. Fulton, R. Fulton, G. Weinstock, R. M. Durbin, J. Burton, D. M. Carter, C. Churcher, A. Coffey, A. Cox, A. Palotie, M. Quail, T. Skelly, J. Stalker, H. P. Swerdlow, D. Turner, A. De Witte, S. Giles, R. A. Gibbs, D. Wheeler, M. Bainbridge, D. Challis, A. Sabo, F. Yu, J. Yu, J. Wang, X. Fang, X. Guo, R. Li, Y. Li, R. Luo, S. Tai, H. Wu, H. Zheng, X. Zheng, Y. Zhou, G. Li, J. Wang, H. Yang, G. T. Marth, E. P. Garrison, W. Huang, A. Indap, D. Kural, W.-P. Lee, W. Fung Leong, A. R. Quinlan, C. Stewart, M. P. Stromberg, A. N. Ward, J. Wu, C. Lee, R. E. Mills, X. Shi, M. J. Daly, M. A. DePristo, D. Altshuler, A. D. Ball, E. Banks, T. Bloom, B. L. Browning, K. Cibulskis, T. J. Fennell, K. V. Garimella, S. R. Grossman, R. E. Handsaker, M. Hanna, C. Hartl, D. B. Jaffe, A. M. Kernytsky, J. M. Korn, H. Li, J. R. Maguire, S. A. McCarroll, A. McKenna, J. C. Nemesh, A. A. Philippakis, R. E. Poplin, A. Price, M. A. Rivas, P. C. Sabeti, S. F. Schaffner, E. Shefler, I. A. Shlyakhter, D. N. Cooper, E. V. Ball, M. Mort, A. D. Phillips, P. D. Stenson, J. Sebat, V. Makarov, K. Ye, S. C. Yoon, C. D. Bustamante, L. Clarke, P. Flicek, F. Cunningham, J. Herrero, S. Keenen, E. Kulesha, R. Leinonen, W. M. McLaren, R. Radhakrishnan, R. E. Smith, V. Zalunin, X. Zheng-Bradley, J. O. Korbel, A. M. Stütz, S. Humphray, M. Bauer, R. Keira Cheetham, T. Cox, M. Eberle, T. James, S. Kahn, L. Murray, A. Chakravarti, K. Ye, F. M. De La Vega, Y. Fu, F. C. L. Hyland, J. M. Manning, S. F. McLaughlin, H. E. Peckham, O. Sakarya, Y. A. Sun, E. F. Tsung, M. A. Batzer, M. K. Konkel, J. A. Walker, R. Sudbrak, M. W. Albrecht, V. S. Amstislavskiy, R. Herwig, D. V. Parkhomchuk, S. T. Sherry, R. Agarwala, H. M. Khouri, A. O. Morgulis, J. E. Paschall, L. D. Phan, K. E. Rotmistrovsky, R. D. Sanders, M. F. Shumway, C. Xiao, G. A. McVean, A. Auton, Z. Iqbal, G. Lunter, J. L. Marchini, L. Moutsianas, S. Myers, A. Tumian, B. Desany, J. Knight, R. Winer, D. W. Craig, S. M. Beckstrom-Sternberg, A. Christoforides, The 1000 Genomes Project Consortium, Corresponding author, Steering committee, Production group: Baylor College of Medicine, BGI-Shenzhen, Broad Institute of MIT and Harvard, Illumina, Life Technologies, Max Planck Institute for Molecular Genetics, Roche Applied Science, Washington University in St Louis, Wellcome Trust Sanger Institute, Analysis group: Agilent Technologies, Baylor College of Medicine, Boston College, Brigham and Women’s Hospital, T. H. G. M. D. Cardiff University, Cold Spring Harbor Laboratory, Cornell and Stanford Universities, European Bioinformatics Institute, European Molecular Biology Laboratory, Johns Hopkins University, Leiden University Medical Center, Louisiana State University, US National Institutes of Health, Oxford University, The Translational Genomics Research Institute, A map of human genome variation from population-scale sequencing. Nature. 467, 1061–1073 (2010).

39. G. Bhatia, N. Patterson, S. Sankararaman, A. L. Price, Estimating and interpreting FST: The impact of rare variants. Genome Res. 23, 1514–1521 (2013).

40. J. K. Pickrell, J. C. Marioni, A. A. Pai, J. F. Degner, B. E. Engelhardt, E. Nkadori, J.-B. Veyrieras, M. Stephens, Y. Gilad, J. K. Pritchard, Understanding mechanisms underlying human gene expression variation with RNA sequencing. Nature. 464, 768–772 (2010).

41. A. Lea, J. Tung, X. Zhou, A flexible, efficient binomial mixed model for identifying differential DNA methylation in bisulfite sequencing data. PLoS Genet. 11, e1005650 (2015).

42. J. Tung, X. Zhou, S. C. Alberts, M. Stephens, Y. Gilad, The genetic architecture of gene expression levels in wild baboons. Elife. 4, 1–22 (2015).

43. C. A. Buerkle, Z. Gompert, Population genomics based on low coverage sequencing: how low should we go? Molecular Ecology. 22, 3028–3035 (2013).

44. W. N. Venables, B. D. Ripley, Modern Applied Statistics with S. Statistics and Computing (2002), doi:10.1007/978-0-387-21706-2.

45. M. Lesnoff, R. Lancelot, aod: Analysis of Overdispersed Data. R package version 1.3. 2012 (2016).

46. R. Kofler, R. V. Pandey, C. Schlötterer, PoPoolation2: identifying differentiation between populations using sequencing of pooled DNA samples (Pool-Seq). Bioinformatics. 27, 3435–3436 (2011).

47. G. dos Santos, A. J. Schroeder, J. L. Goodman, V. B. Strelets, M. A. Crosby, J. Thurmond, D. B. Emmert, W. M. Gelbart, FlyBase Consortium, FlyBase: introduction of the Drosophila melanogaster Release 6 reference genome assembly and large-scale migration of genome annotations. Nucleic Acids Res. 43, D690–7 (2015).

48. M. Wang, Y. Zhao, B. Zhang, Efficient Test and Visualization of Multi-Set Intersections. Scientific Reports. 5, 16923 (2015).

49. R. Kofler, C. Schlötterer, Gowinda: unbiased analysis of gene set enrichment for genome-wide association studies. Bioinformatics. 28, 2084–2085 (2012).

50. G. F. Berriz, J. E. Beaver, C. Cenik, M. Tasan, F. P. Roth, Next generation software for functional trend analysis. Bioinformatics. 25, 3043–3044 (2009).

51. J. Thurmond, J. L. Goodman, V. B. Strelets, H. Attrill, L. S. Gramates, S. J. Marygold, B. B. Matthews, G. Millburn, G. Antonazzo, V. Trovisco, T. C. Kaufman, B. R. Calvi, FlyBase Consortium, FlyBase 2.0: the next generation. Nucleic Acids Res. 47, D759–D765 (2019).

52. A. M. Taverner, L. J. Blaine, P. Andolfatto, bioRxiv, in press, doi:10.1101/2020.08.05.237594.

53. G. S. C. Slater, E. Birney, Automated generation of heuristics for biological sequence comparison. BMC Bioinformatics. 6, 31 (2005).

54. J. B. Lack, J. D. Lange, A. D. Tang, R. B. Corbett-Detig, J. E. Pool, A Thousand Fly Genomes: An Expanded Drosophila Genome Nexus. Mol. Biol. Evol. 33, 3308–3313 (2016).

55. A. Löytynoja, N. Goldman, A model of evolution and structure for multiple sequence alignment. Philos Trans R Soc Lond B Biol Sci. 363, 3913–3919 (2008).

56. S. F. Altschul, W. Gish, W. Miller, E. W. Myers, D. J. Lipman, Basic local alignment search tool. Journal of Molecular Biology. 215, 403–410 (1990).

57. R. C. Edgar, MUSCLE: a multiple sequence alignment method with reduced time and space complexity. BMC Bioinformatics. 5, 113 (2004).

58. A. J. Lea, D. Martins, J. Kamau, M. Gurven, J. F. Ayroles, Urbanization and market integration have strong, nonlinear effects on cardiometabolic health in the Turkana. Science Advances. 6, eabb1430 (2020).

59. A. J. Lea, J. Altmann, S. C. Alberts, J. Tung, Developmental Constraints in a Wild Primate. Am Nat. 185, 809–821 (2015).

