## Supplementary material for "Diet unmasks genetic variants that regulate lifespan in outbred *Drosophila*": Methods and Supplementary Figures

**This PDF file includes:**

Materials and Methods

Figs. S1 to S9

Tables S1 to S12 are provided in a separate spreadsheet

**Materials and Methods**

S1. Fly population

600 isofemale lines were established from wild *Drosophila melanogaster* flies collected in Princeton, NJ in March-April 2017. Five males and five females per line were placed in a medium-sized population cage (W32.5 x D32.5 x H32.5 cm) and allowed to mate freely. After 15 generations the flies were split into two large cages (W47.5 x D47.5 x H47.5 cm) to increase the population size. The outbred populations were kept in a regime of 25°C, 65% humidity, and a 12h/12h light/dark cycle. They were kept on media that we refer to as "control" diet, which had the following composition: 0.6% agar, 0.7% potassium sodium tartrate, 0.1% $CaCl_2$ 2H2O, 2.5% yeast, 6.1% cornmeal, 0.6% propionic acid, 2.5% sucrose and 5% glucose. At the time of our experiments, flies had been randomly mating and maintained on the control diet for approximately 25 generations.

S2. Fly husbandry and experimental design

*Main experimental setup*

Over a period of four days, bottles with fresh control media were placed in the two cages to collect eggs from adult flies. During this time, we performed six rounds of egg laying for each of the cages, with each egg lay lasting 7-20 hours to maintain a roughly constant egg density, and thus, limit extreme differences in larval density.

Flies that eclosed 4-6 days after the first fly eclosed were collected and randomly assigned to two pools, one consisting of flies from egg lay 1, 2, and 4, and the second from egg lay 5 and 6. We refer to these two pools in the main text as A and B. Flies that eclosed very early (day 1-3 after the first fly eclosed) or very late (day 7-10 after the first fly eclosed) were not included in the experiment to control for the possibility that extremely fast or slow development may influence longevity.

Approximately 1000 flies were randomly sampled from each of the two pools (A and B) using weak vacuum aspiration and stored at -80C in 50mL Falcon tubes. We refer to these pools as "$T_0$" flies as they represent the starting point of the experiment when flies were 2±1 days old. We note that the minor allele frequency spectrum at $T_0$ was very similar between pool A and B flies (Figure 1C).

The rest of the flies were randomly allocated into six replicate cages in the following way: groups of ~10,000 flies from pool A (egg lay 1, 2, and 4) were placed in four large cages, and groups of ~10,000 flies from pool B (egg lay 5 and 6) were placed in two large cages, for a total of six cages. To evaluate the effect of different environments on fly longevity, three cages were fed with control media (7.5% sugar) and the other three with "high sugar" media (22.5% sugar). High sugar media had the same composition as control media with an additional 5% glucose and 10% sucrose. The cages were maintained on a given diet for the duration of the experiment.

Food was freshly provided in petri dishes every 3-4 days, and old dishes were discarded to ensure that larvae did not develop into adults. In addition, every 3-7 days ~500 live flies were sampled from a given cage using weak vacuum aspiration and stored in 1.5mL Eppendorf tubes at -80C; we refer to these samples as "intermediate time points". When the last ~500 surviving flies were left in a given cage, all flies were collected and stored in 1.5mL Eppendorf tubes at -80C; we refer to these samples as "$T_N$" flies. At the $T_N$ time point, flies were between 32-59 days old depending on the cage (see Table S1-S2 for cage-specific sample collection schedules).

Throughout the experiment, all cages were kept in a single 25°C incubator at 65% relative humidity with a 12h/12h light/dark cycle.

*Follow up experiment to validate sex-by-environment effects on longevity*

To validate the sex-by-diet effects on longevity that we observed in the main experiment, we performed a vial-based survival assay where the lifespan of individual flies was quantified. Eggs were collected from the original outbred population, and flies that eclosed 4-6 days after the first fly eclosed were sexed and split across 42 vials each with 5mL of media and the following design: (i) 6 replicates, each containing 10 males on a control diet, (ii) 6 replicates, each containing 10 females on a control diet, (iii) 9 replicates, each containing 5 males and 5 females on a control diet, and (iv-vi) the same replicate structure for the high sugar diet. We designed this experiment to include flies housed in single sex or mixed sex groups, in order to evaluate whether the sex-by-diet interactions we observed in the main experiment were driven by interactions between the sexes (e.g., mating). Flies were transferred to fresh media every 3-4 days, and the number and sex of dead flies was recorded at each transfer. All vials were kept in a single 25°C incubator at 65% relative humidity with a 12/12 hour light/dark cycle.

To estimate the effect of sex and diet on lifespan, we fit mixed effects Cox proportional hazards models in the R package *coxme* (*29*). Specifically, for the male-only and female-only vials we fit models with a fixed effect of diet and a random effect of vial. For the mixed sex vials, we fit models with fixed effects of sex and diet and a random effect of vial (see results presented in Table S5). The Kaplan-Meier survival curves presented in Fig. S2 were created with the R package *survival* (*30*, *31*).

*Follow up experiment to validate effects of candidate genes on survival*

We selected two candidate genes to validate their predicted effects on lifespan, *lovit* and *midway.* To test the effect of each gene on lifespan, but in particular to test whether the lifespan effect is modulated by diet, we compared the survival of loss-of-function mutant flies to that of four control lines on both a control and high sugar diet. Five males and five females were placed together in vials with 5mL of media and the number of sex-specific deaths was recorded every three days when the flies were transferred to fresh food vials. We used six replicates per group, for a total of 60 flies per line per diet. This experiment was conducted at home under uncontrolled temperature and humidity conditions, because Princeton University was closed for research as a result of the COVID-19 pandemic. The replicate vials were split between two houses and therefore our results are robust to variation in temperature and humidity.

Loss-of-function lines were obtained from Bloomington Drosophila Stock Center: *lovit* (FBgn0267429, BDSC_83374) and *midway* (FBgn0004797, BDSC_5095). Although all mutant lines were reported in the literature as homozygous viable, very few flies from the *midway* line were indeed viable homozygous and therefore this gene was tested in its heterozygous state. The control lines were: DGRP_439, DGRP_181, Canton-S, and yw.

To estimate the effects of genotype, diet, and the interaction between genotype and diet on lifespan, we fit mixed effects Cox proportional hazards models in the R package *coxme* (*29*). We analyzed data for each of the candidate genes separately, with genotype coded as "mutant" for the loss of function line and a category called "control" which included all 4 of the control lines. All models included a random effect of vial. For each candidate gene, we also explored a more complicated model that included sex and the interaction between sex and diet, sex and genotype, and a three-way interaction between sex, diet, and genotype. Full results from these models are presented in Table S10. The Kaplan-Meier survival curves presented in Figures 3-4 in the main text were created with the R package *survival* (*30*, *31*).

S3. DNA samples

*DNA extraction, library preparation, and sequencing*

We used a low-coverage DNA sequencing approach to estimate allele frequencies in each of the six replicate cages from the main experiment, for the $T_0$ and $T_N$ samples as well as the intermediate time points samples. Each previously frozen fly from $T_0$ and $T_N$ time points was placed in a well of a 96-well plate for DNA extraction, and the DNA sequencing data was used to sex each individual (see *Processing of DNA sequencing data to determine sex*). Due to the strong bias in sex-specific survival between $T_0$ and $T_N$, the flies from the intermediate time points were sexed on dry ice previous to DNA extraction to ensure that both sexes were sequenced.

A given plate was always filled with flies from the same time point and cage, with three random wells left empty to serve as negative controls. One 2.8mm stainless steel grinding bead (OPS diagnostics, #089- 5000-11) and 100µl of lysis buffer were added to each well. Flies were homogenized for 10 minutes at maximum speed in a Talboys High Throughput Homogenizer (#930145). The resulting lysate was transferred to a new 96-well plate for DNA extraction, which was performed using a Multi-Well Plate Vacuum Manifold (#5017) and the Acroprep advance 1mL DNA binding plates (#8132), both from Pall Life Sciences.

After DNA extraction, library preparation was performed using a CyBio® FeliX liquid handling robot (Analitik Jena) to allow for high-throughput while reducing the variability introduced by manual handling of individual samples. Library preparation generally followed the strategy outlined in (*32*), using the oligo sequences and homemade Tn5 recipe therein. Specifically, 10µl (100µM) forward oligo adapter A and 10µl (100µM) reverse oligo (Tn5MErev) were mixed with 80µl reassociation buffer (10mM Tris pH 8.0, 50mM NaCl, 1mM EDTA), and annealed in a thermocycler following the program: 95°C 10min, 90°C 1min, reduce temperature by 1°C/cycle for 60 cycles, hold at 4°C. The same procedure was also performed using forward oligo adapter B. To load the pre-annealed adapters onto the Tn5, 45µl of Tn5 was combined with 9µl of pre-annealed adapter A and 9µl of pre-annealed adapter B (both at 10µM) and incubated in a thermal cycler for 30min at 37°C. The pre-charged Tn5 was then diluted with reassociation buffer:glycerol (1:1), in a ratio of 1 part reassociation buffer:glycerol to 1 part pre-charged Tn5. 2µl of each DNA sample was then mixed with 1µl of precharged Tn5, 2µl of 5X TAPS buffer pH 8.5 (50mM TAPS, 25mM MgCl2, 50% v/v DMF), and 5µl of water. The solution was incubated for 7min at 55°C, after which 2.5µl of 0.2% SDS (Promega, #V6551) was added and incubated in a thermal cycler for 7min at 55°C to dissociate the Tn5 from the DNA. Final library amplification was then performed by combining 2µl of the tagmentation reaction with 7µl of OneTaq HS Quick-Load 2x (NEB, #M0486L), 1µl of the i5 primer (5uM), 1µl of the i7 primer (5µM), and 4µl of water. This mixture was amplified on a thermal cycler with the following program: 68°C 3min, 95°C 30sec, [95°C 10sec, 55°C 30sec, 68°C 30sec] for 18 cycles, 68°C 5min.

For each 96 well plate, 2μl of the final PCR amplification reactions were pooled (for a total volume of 192μl) and size selected using a double-sided Agencourt AMPure XP bead (Beckman Coulter) cleanup approach for an average insert size of 400bp. The ratio of beads to sample for the first and second cleanups were 0.6x and 0.4x, respectively. Pooled and cleaned libraries were visualized on an Agilent TapeStation, and sequenced on the Illumina NovaSeq S4 platform at the New York Genome Center using 100 bp PE reads (see Fig. S1 for distributions of sample-specific read depths). DNA-seq libraries were sequenced in 3 batches, with two batches containing libraries constructed from $T_0$ and $T_N$ flies, and the final batch containing libraries constructed from flies collected at intermediate time points (see Table S2 for sample sizes).

Protocols and subroutines for library preparation using the CyBio® FeliX liquid handling robot are available upon request.

*Processing of DNA sequencing data*

Following sequencing, we removed adapter contamination and low-quality bases from each library using *cutadapt* (*33*). We mapped the trimmed reads to the *Drosophila melanogaster* reference genome (v6.23) using *BWA* (*34*), and retained only uniquely mapped reads. Next, we removed duplicate reads generated during PCR using *Picard* (http://broadinstitute.github.io/picard), and performed base quality score recalibration, application, and individual variant calling using the Genome Analysis Toolkit (v3.2-2) (*35*). Because no validated reference set of genetic variants were available for the outbred population we used for our experiments, we relied on an iterative approach for base quality score recalibration. Specifically, we performed an initial round of base quality score recalibration, individual variant calling (using HaplotypeCaller), and joint genotyping (using GenotypeGVCFs) on 550 samples from the first batch of sequencing with high read depths (>1.5 million PE reads per individual). We then used GATK's VariantFiltration to perform hard filtering and construct a set of high confidence variants with quality scores ≥100 that passed all filters for variant confidence (variants failed if QD < 2.0), mapping quality (variants failed if MQ < 35.0), strand bias (variants failed if FS > 60.0), mapping quality (variants failed if MQRankSum < −12.5) and read position bias (variants failed if ReadPosRankSum < −8.0). We also removed variants that were not genotyped in >10% of samples. We used this high confidence set of "known sites" in a second round of base quality score recalibration for all samples (including the original high-coverage set of 550). Following the second round of base quality score recalibration, joint genotyping was performed in batches (using GenotypeGVCFs) for all samples from a given time point and/or replicate cage.

Our main analyses focused on comparisons between the $T_0$ and $T_N$ time points. Therefore, we first filtered for sites with non-zero read counts in >25% of samples from all six $T_N$ cages as well as the two $T_0$ cages. We also filtered for sites with quality scores >20, a Hardy Weinberg p-value >$10^{-6}$, and a minor allele frequency >5% in all $T_0$ and $T_N$ cages. This filtering was performed using *bcftools* (http://samtools.github.io/bcftools/bcftools.html) and resulted in a set of 311,180 SNPs. Next, we used *Plink* to perform LD filtering such that in a moving window of 500 SNPs (with a 50 SNP offset), the pairwise r2 between any two SNPs was always < 0.5 (*36*). This step removed 14.6% of SNPs. Finally, we removed individual samples with non-zero read counts for <10% of sites passing filters (see Table S2 for final sample sizes) and we removed 290 sites with unusually high mean read depths (>10x) across the filtered sample set. This set of filtering criteria left us with 271,246 sites for analyses. For those sites, we extracted the number of reads that mapped to the alternate and reference alleles for each individual sample using *bcftools* (http://samtools.github.io/bcftools/bcftools.html).

*Processing of DNA sequencing data to determine sex*

Flies were sexed from the sequencing data based on the ratio of the number of reads that mapped uniquely to the X chromosome and the number of reads that mapped uniquely to all of the autosomes. This ratio followed a clear binary distribution (Fig. S8), and we considered flies with ratios < 0.13 to be male and flies with ratios between 0.18 and 0.25 to be female.

S4. Population genetic analyses

To characterize the genetic diversity of pool A and B at $T_0$, as well as each of the 6 cages at $T_N$, we used the Genome Analysis Toolkit (v3.8) (*35*) to output variant and invariant sites using the function GenotypeGVCFs with the flag -includeNonVariantSites. Each chromosome arm was processed separately, and cages with more than 500 individuals were divided into two groups to minimize computational footprint but were merged again for the estimation of the different diversity parameters.

For each site, reads generated from each individual were used to assign that individual to the "reference" or "alternate" genotype. If an individual had only one read for a given site, it was considered homozygous for the genotype called from this read (i.e., contributing two reads to such genotype). Only monoallelic and biallelic sites were used in further analysis.

Using custom scripts, we calculated site level and population level diversity parameters. Per-site polymorphism (pi) was calculated as (*N_alleles*/(*N_alleles*-1))*2**p***q*, where *N_alleles* is the total number of genotyped alleles in the population, *p* is the frequency of the major allele, and *q* is the frequency of the minor allele. Population pi was calculated as the average of the per-site polymorphism. Number of segregating sites in the population was calculated as the total number of sites that are biallelic. The folded allele frequency spectrum was calculated as the number of sites with a given minor allele frequency, in bins of 0.05 (ranging from 0 to 0.495). For the calculation of the fixation index (Fst) between pair of cages, only sites that remained monoallelic or biallelic after combining the read counts of both cages were used. Fst was then calculated as (*pi_T*-*pi_S*)/*pi_T*, where *pi_T* is per-site pi across the two cages and *pi_S* is the average of the two cage-specific per-site pi. We then calculated Fst at the population level using two approaches, the ratio of averages (*37*) and the average of ratios (*38*), where the ratio of averages is generally higher than the average of ratios (*39*). For the ratio of averages estimate, we calculated the average *pi_T* and *pi_S* across all sites, and such averages were used in the Fst calculation. For the average of ratios estimate, we first calculated Fst for each site, and then the average across all site-specific Fst. The above metrics, calculated for each cage, are presented in Table S3.

S5. <u>Comparison of two statistical approaches to test for allele frequency changes with age</u>

*Overview of approaches*

Our main goal was to understand how allele frequencies differed between the $T_0$ and $T_N$ time points in each of the dietary environments. The most commonly used test to compare allele frequencies between two sample groups (e.g., treatments or time points) across multiple replicates is the Cochran–Mantel–Haenszel (CMH) test. This test can either be performed using counts of the number of individuals carrying a given allele in each replicate (when per-individual genotype data are available), or using counts of the number of reads supporting a given allele in each replicate (when pool-seq data are generated).

Given that we used individually barcoded libraries, our data consists of the number of reads that mapped to the alternate versus reference allele for each individual fly. As a result, we can implement the CMH test by creating perfectly weighted pool-seq data *in silico* where each fly contributes exactly one read to the estimation of allele counts. In this way, we are able to control for the biases in allele count estimation driven by unequal representation of individual samples. However, this approach has two main limitations: first, by downsampling to one read per sample the power to detect allele frequency differences is reduced because sample size corresponds to the number of individuals with sequencing data and not to the total amount of sequencing reads in the sample. Second, the CMH test does not allow the user to control for covariates (e.g., sex or batch) or to compare multiple sample groups (e.g., high sugar and control diets) within the same statistical test.

To overcome such limitations, we considered a beta-binomial regression framework to directly model individual-specific read count data and to control for covariates. Because we used a low-coverage (~1-2x) sequencing approach, our data are not appropriate for assigning individual genotypes, however beta-binomial regressions offer an alternative for genomic data types such as ours where the outcome of interest is per-individual reads supporting one state / total reads generated (as in analyses of allele specific expression (*40*, *41*) or bisulfite sequencing data (*42*)). In our case, we modeled the number of reads that mapped to the alternate allele / total reads generated for each individual; this approach to low coverage genotype data was also proposed by (*43*).

*Simulations*

To estimate differences in statistical power between the CMH and beta-binomial approaches, we simulated read counts (reads mapped to alternate allele / total reads generated) for 1000 loci and 2000 individuals (n=1000 at $T_0$ and two replicates of n=500 at $T_N$). We performed 11 replicate simulations that were identical in sample size and in the number of loci, but varied in the percent decrease of the alternate allele between $T_0$ and $T_N$ (ranging from 0% to 20%).

Specifically, for a given replicate simulation, we simulated the allele frequencies at $T_0$ for 1000 loci by randomly sampling from our real dataset (using allele frequencies for the set of all sites passing filters). For each locus, we then decreased the $T_0$ allele frequency by 0-20%

depending on the simulation, and set this value as the $T_N$ allele frequency. Finally, we simulated genotypes for $T_0$ and $T_N$ flies by drawing from a binomial distribution parameterized by the sample size and the sample allele frequency.

To translate these simulated genotypes into count data obtained from low coverage sequencing, we first simulated total read counts $r_i$ for each individual $i$ by sampling from a negative binomial distribution: $r_i \sim NB(t,p)$. Here, $t$ and $p$ are site specific parameters estimated from the real sequencing data. Specifically, we generated 5000 sets of $t$ and $p$ parameters by fitting a negative binomial distribution to the total read count data from randomly selected sites that passed filters in our real dataset (using the function *fitdistr* in the R package *MASS* (*44*)). To simulate total read counts for a given site, we randomly selected one of these parameter sets to produce the total number of reads. Finally, we simulated the number of alternate allele reads for each individual at that locus by drawing from a binomial distribution parameterized by the number of total reads ($r_i$) and the focal individual's genotype which was simulated previously and coded here as 0, 0.5, or 1 (for homozygous reference, heterozygous, and homozygous alternate respectively).

*Analysis of simulated data*

Simulated count data were analyzed with either the CMH or beta-binomial approach described above, and power was determined as the proportion of simulated loci with changes in allele frequency between time points detected at a p-value threshold of $10^{-6}$.

Beta-binomial regressions testing for a fixed effect of time were implemented with the R package *aod* (*45*). Specifically, the number of reads for the alternate allele / total reads at each site was modelled as a function of time for each simulated dataset.

For the CMH implementation, we used a down-sampling strategy to ensure that only one read per individual was used in the estimation of allele counts per cage. First, the individuals from $T_0$ were randomly assigned to two $T_0$ replicates to fulfill the CMH assumption of independence and to be able to run the test using two replicates of the $T_N$-$T_0$ comparison. Then, for each site per replicate cage, we sampled one read per individual and calculated the total counts for the reference and alternate allele. We repeated this procedure 1000 times and calculated the average reference and alternate counts across all 1000 rounds to the closest even integer. The average reference and alternate counts for each replicate cage were used in the CMH test implemented in *PoPoolation2* (*46*) using the following parameters: --min-count=12, --min-coverage=50, --max-coverage=1000, --population= $T_{N\_1}$ - $T_{0\_1,}$ $T_{N\_2}$ - $T_{0\_2}$. An independent CMH test was run for each one of the simulated datasets with allele frequency differences between $T_N$ and $T_0$ ranging from 0% to 20%. When we compared the results of the two statistical approaches, we found that the beta-binomial provided more power to detect allele frequency differences, as expected (Fig. S9).

S6. <u>Identification of loci associated with variation in lifespan</u>

*Comparisons between $T_0$ and $T_N$ to identify genetic effects on longevity*

Our simulations revealed that a beta-binomial approach should have greater power to detect changes in allele frequency relative to a CMH test, and we therefore implemented beta-binomial regressions as the main statistical approach to analyze the real data. For each individual *i*, we modeled the number of reads mapped to the alternate allele and the total read counts at each locus as:

$$y_i = Bin(r_i, \pi_i),$$

where $r_i$ is the total read count for the *i*th individual, $y_i$ is the number of reads mapped to the alternate allele for that individual and $\pi_i$ is an unknown parameter that represents the genotype of individual *i* at the site (where 0=homozygous reference, 0.5=heterozygote, and 1=homozygous alternate). We then used a logit link implemented in a beta-binomial regression to model $\pi_i$ as a function of several parameters:

$$log(\pi_i/(1-\pi_i)) = \mu + p_i\beta_p + b_i\beta_b + s_i\beta_s + c_i\beta_c + h_i\beta_h + \varepsilon_i$$

where $\mu$ is the intercept, $\varepsilon_i$is residual model error, $p_i$ is the starting population pool (A or B; see the *Main experimental setup* section of Text S2), $b_i$ is the sequencing batch (1 or 2), $s_i$ is sex (male or female) and a 3-level factor representing time and condition ($T_0$, $T_N$ high sugar, and $T_N$ control) configured with two contrasts: $T_0$ versus $T_N$ high sugar (represented by $\beta_h$) and $T_0$ versus $T_N$ control (represented by $\beta_c$). It is important to note that we were unable to explicitly model an interaction between time and diet, because the $T_0$ sample is common to both diets. We considered sites to have a time effect specific to the high sugar environment if the p-value associated with $\beta_h$ passed a 10% FDR and the nominal p-value associated with $\beta_c$ was >0.05 (and vice versa to identify sites with a time effect specific to the control environment). We considered sites to have a time effect shared between dietary conditions if the p-value associated with $\beta_c$ or $\beta_h$ passed a 10% FDR in at least one environment and the nominal p-value in the other environment was <0.05. Q-values for each site were empirically derived by permuting the 3-level factor representing time and condition 10 times, and rerunning our analyses. We then calculated empirical q-values for each site (for $\beta_c$ and $\beta_h$) using the null distributions obtained via permutation and the R function available here: https://rdrr.io/github/janderson94/Revolution/man/perm.fdr.html.

We also implemented the CMH approach using our real data, and using the same down sampling approach described in the previous section (i.e., one read, per site, per fly). Fig. S11 shows that the correlation between p-values obtained from the CMH versus beta-binomial approaches are very similar. Further, we found that 96% and 92% of sites that we identified with the CMH test as having a significant genetic effect on longevity in the high sugar and control environments, respectively, were also detected by the beta-binomial approach. However, as expected from our simulations, the absolute number of significant sites detected by a CMH approach was much lower (specifically, 5.42-fold lower).

*Evaluating definitions of GxE versus shared genetic effects*

As noted in the above section, we used a combination of p-value and FDR thresholds to define “shared” and “GxE” SNPs. Specifically, we consider shared SNPs to be those that pass a 10% FDR in one environment and a nominal p-value threshold of 0.05 in the other; we consider GxE SNPs to be those that pass a 10% FDR in one environment and fail to reach a nominal p-value threshold of 0.05 in the other. This reciprocal thresholding could potentially lead to truly shared SNPs being falsely defined as GxE if they fail to replicate in the second environment.

To understand how often this might occur, we used permutations and simulations. First, we permuted the labels for $T_N$ HS and $T_N$ CTRL within the 3-way factor representing time and condition 10 times, and reran our analyses. In theory, this permutation scheme should eliminate our ability to detect sites with true GxE effects but not shared effects, since the dietary labels (HS vs CTRL) are scrambled but the time labels ($T_N$ vs $T_0$) are preserved. We then applied the same procedure we used to identify GxE effects in the real data to the permuted data; specifically, we empirically derived q-values using a Storey-Tibshirani approach and defined GxE as sites that pass a 10% FDR in one environment and fail to reach a nominal p-value threshold of 0.05. We recorded the proportion of total sites tested that we assigned as GxE in the permuted data and compared this to the proportion of sites tested that we assigned as GxE in the real data. We estimate that we assign 8.83x more sites as GxE in the real data relative to the permuted data (equivalent to a false positive rate of 11.3%). To explore the robustness and behavior of this false positive rate estimate, we also repeated the above analyses but varied the FDR threshold used to call a site significant in one environment; these results are presented in Figure S5C.

Second, we ran simulations similar to those described in the *Simulations* section of Text S6. Specifically, we simulated read count data (reads mapped to alternate allele / total reads generated) for 1000 loci and 3000 individuals (n=1000 at $T_0$ and n=1000 at $T_N$ for two environments). We performed 121 replicate simulations that were identical in sample size and in the number of loci, but varied in the percent decrease of the alternate allele between $T_0$ and $T_N$ in each of the two environments. In particular, we explored a grid where the alternate allele decreased between 0% and 20% in environment 1 and environment 2 (see x- and y-axes of Fig. S6). For each simulation, we analyzed the data with a beta-binomial model and extracted the p-values associated with the time effect in environment 1 and environment 2. We then asked what proportion of loci were classified as GxE with larger effects in environment 2, using the following definition: $p<10^{-4.5}$ (similar to the 10% FDR threshold in the real data) in environment 2 and $p>0.05$ in environment 1. We also asked what proportion of loci were classified as shared using the following definition: $p<10^{-4.5}$ in one environment and $p<0.05$ in the other environment.

The results of these simulations are presented in Fig. S6A-B and suggest a few key take home points. First, we are more likely to classify a site as GxE if there are large differences in effect sizes between the two environments (relative to sites with only modest differences in effect sizes). Second, we do not misclassify sites with equal effect sizes between environments as GxE (see the diagonal of Fig. S6B). Third, our set of shared sites may include a small proportion of sites that are technically GxE, in that they exhibit large and similar (but not identical) effect sizes between the two environments (see the off diagonal of Fig. S6A).

S7. Analyses of lifespan-associated SNPs and genes

*Testing for enrichment of lifespan-associated SNPs in genomic features*

Lifespan-associated SNPs were annotated as falling in introns, exons, UTRs, or intergenic regions based on the genome annotation file for the *D. melanogaster* reference genome v6.23 from FlyBase (*47*) (ftp://ftp.flybase.net/genomes/Drosophila_melanogaster/dmel_r6.23_FB2018_04/gtf/). Using a hypergeometric test and the background set of all tested SNPs, we asked whether lifespan-associated SNPs were enriched for any particular genomic feature. We performed this analysis separately for SNPs with shared effects and SNPs with GxE effects (i.e., genetic effects that were stronger in one environment relative to the other). These results are provided in Table S7.

*Testing for overlap with previous studies*

We assigned each of our lifespan-associated SNPs to a given gene if they fell within the gene body or within 1kb of the gene's annotated TSS or TES. Using the list of genes near SNPs with significant shared or GxE effects, we compared our gene-level results with six previous studies assessing the genetic basis of lifespan variation in *D. melanogaster* (*14*, *18*, *19*, *25*–*27*), and with the database *GenAge* which harbors a manually curated list of aging related genes in *D. melanogaster* (*28*). We tested for pairwise overlap between each gene list (Table S8) using the *SuperExactTest* package in R (*48*). As the background set, we used the 15,751 genes that were tagged by any tested SNP in our analysis. Significance was determined at 1% FDR threshold, and full results are presented in Table S9.

*GO enrichment analysis*

To identify whether the set of genes with lifespan-associated SNPs was enriched for any biological process or function we used *Gowinda* (*49*). This program accounts for the biases introduced by gene length and gene clustering by randomly sampling gene sets from the complete list of SNPs used in our analysis. *Gowinda* was run in "gene" mode, using 100,000 permutations, only considering GO terms that contain a minimum of three genes, and the SNP-to-gene mapping was done by extending 1kb upstream and downstream of the TES and TSS. This analysis was run independently for the two SNP sets: SNPs with shared lifespan effects and SNPs with GxE effects (i.e., effects that were stronger in one environment). The GO information was obtained from *FuncAssociate2* (*50*).

S8. Testing different theories of aging by assigning alleles to particular trajectories

Our main analyses focused on allele frequency comparisons between $T_0$ versus $T_N$; however, we were also interested in the trajectory of allele frequency change for lifespan-associated alleles, as the shape of the trajectory could provide insight into different theories of aging. In particular, we were interested in understanding whether lifespan-associated alleles tended to fit one of two patterns: (i) an allele is maintained at a constant frequency up to a given age but then steadily declines, suggesting the allele only affects longevity in late life as predicted by the mutation accumulation theory of aging (*4*); or (ii) an allele decreases in frequency early in life but increases again in late life, indicating trade-offs as proposed by the antagonistic pleiotropy theory of aging (*5*). We also considered a third possible pattern, in which an allele decreases at a constant rate across all ages suggesting consistent longevity effects throughout the lifespan (Figure 5A). We note that this pattern was unexpected and serves as somewhat of a null model, given that it does not fit any known evolutionary theories of aging.

To test different theories of aging by assigning alleles to particular patterns of change across the lifespan, we focused on the set of 2246 SNPs that had significant effects on longevity in at least one environment. Using data from either the $T_0$ plus high sugar samples or the $T_0$ plus control samples alone, we fit three beta-binomial models (described below) to data from all $T_0$ and $T_N$ samples, as well as all intermediate time points samples. For the high sugar dataset, we focused on 2243 SNPs with significant shared or high sugar-biased effects on longevity; for the control dataset, we focused on 1545 SNPs with significant shared or control-biased effects on longevity. We analyzed the high sugar and control datasets separately because the set of possible ages in each environment was very different (age range for high sugar samples: 0-33 days; age range for control samples: 0-58 days; Table S1).

For each dataset and SNP set, we modeled the number of reads mapped to the alternate allele / total read counts at each locus as a function of the following main effects: starting population pool, sequencing batch, sex, and age. Age was coded as a numeric variable and modeled linearly (to test the constant/linear model), quadratically (to test the antagonistic pleiotropy theory), or linearly after coding all early time points as the same middle-aged value (to test the mutation accumulation theory). For the mutation accumulation theory model, we tried various definitions of “early time points”, ranging from <36 to <51 days for control samples and

<21 to <27 days for high sugar samples (these ranges were designed to search the second half of the fly lifespan on each diet for breakpoints in allele frequency dynamics). We then chose the model with the lowest AIC and used this for each SNP going forward.

To compare all 3 potential trajectories, we extracted the AIC value from each of the three tested models (linear, antagonistic pleiotropy, and the best mutation accumulation model). We then calculated ΔAIC as the difference in AIC between the best model (lowest AIC) and the second-best model (second lowest AIC). To ask whether the ΔAIC value we obtained for a given SNP was larger than expected by chance, we reran our analyses after permuting age for the intermediate time point samples. We left age unpermuted for $T_0$ and $T_N$ samples, because we are assessing the significance of allele frequency dynamics with age not age effects on allele frequencies (only loci known to change in allele frequency between $T_0$ and $T_N$ were included in these analyses in the first place). We considered a site to be confidently assigned to a given trajectory/model if that model had a ΔAIC >97.5% percentile of ΔAIC values recorded across 1000 permuted datasets. Information about SNPs that were confidently assigned to a given trajectory/model is provided in Table S11.

S9. Testing whether derived alleles exhibit a bias toward shortening lifespan

*Identifying the ancestral and derived allele at each locus*

We attempted to identify the ancestral allele at each locus with significant effects on longevity as well as at 10,000 randomly drawn non-significant (FDR>10%) sites using alignments between *D. melanogaster* r6.11 (*51*), *D. simulans* w501, and *D. yakuba* Tai18E2 (Reilly and Deitz *et al.*, in prep). We used two separate approaches to identify the ancestral allele, one focused on coding regions only and one focused on the whole genome.

In the “coding regions" approach, we focused on the longest transcript for each gene. To generate multispecies coding sequence (CDS) alignment between *D. yakuba*, *D. simulans*, and *D. melanogaster*, we first identified one-to-one protein orthologues between the reference genomes of all 3 species using the reciprocal best exonerate approach described in Taverner *et al.* (*52*), with exonerate version 2.4.0 (*53*). To be able to determine the allele frequency of lifespan-associated SNPs from the Nexus world-wide collection of *D. melanogaster* genomes that uses genome assembly r5 (*54*), we extracted the CDS from *D. melanogaster* r5 for each one-to-one ortholog identified in the previous step. Then, using *PRANK* version 170427 (*55*) we aligned the *D. melanogaster* CDS to the sequences from *D. simulans* and *D. yakuba* previously identified as orthologous by exonerate. The final multispecies alignment contains the protein sequence of the longest transcript for 11,820 genes derived from the three reference genomes and 821 *D. melanogaster* individuals from the *Drosophila* Nexus in *D. melanogaster* r5 coordinate space. To convert the genome coordinate of the lifespan-associated SNPs to transcript coordinates, we used the fbgn_fbtr_fbpp_fb_2018_04.tsv conversion file from FlyBase to assign the Fbtr identifier to every SNP. Out of the 4294 lifespan-associated SNPs, 3599 SNPs (84%) were found in the longest transcript of their respective genes with CDS alignment available. Then, to identify the SNP location (r6.23) in the coordinate space of the CDS alignment (r5), we first extracted 25bp on either side of the SNP from the *D.melanogaster* reference genome r6.23. The resulting 51bp sequence was then mapped onto the respective transcript of *D.melanogaster* r5 using *BLAST* version 2.4.0+ (*56*).

For the “whole genome” approach we included SNPs in noncoding as well as coding regions of the genome. A 100 bp window surrounding each lifespan-associated SNP in the *D. melanogaster* reference genome r6.27 was mapped to the *D. simulans* w501 and *D. yakuba* Tai18E2 reference genomes (Reilly and Deitz *et al.*, in prep) using *BLASTN* version 2.9.0 (*56*). Unambiguously mapped regions were then extracted from the *D. simulans* and *D. yakuba* reference genomes and a multi-sequence alignment of the three species was created for each region using *MUSCLE* version 3.8.31 (*57*).

In both approaches, the ancestral state at each SNP was assigned using parsimony; in other words, it was defined as the state shared by at least two of the three species. If all three species showed different states at the focal SNP, the ancestral state was deemed ambiguous.

*Estimating the bias toward the derived allele decreasing lifespan*

After running both the coding regions and whole genome approaches, we decided to focus our analyses on annotations using the "whole genome" approach. We made this decision because we were able to assign ancestral state for a larger set of SNPs using this approach, and because it is not restricted to coding regions that likely have different evolutionary dynamics than the rest of the genome. Specifically, we assigned ancestral state to 1007 lifespan-associated SNPs and 6054 non-significant SNPs using the whole genome approach. Of these 7061 sites, only 737 were also assigned by the coding regions approach, but importantly, sites called by both methods agreed 89% of the time. All ancestral state calls are provided in Table S12.

For SNPs with environmentally shared effects on longevity as well as GxE effects, we estimated the proportion of loci for which the derived allele decreased over time in a given environment. For the set of shared SNPs, we estimated the proportion of loci for which the derived allele decreased over time in each of the two environments separately (but found that these were almost identical). For the set of SNPs with stronger effects in the HS environment, we estimated this same bias in the HS environment only.

To determine whether the bias we observed in the real data was more than expected by chance, we asked how often the derived allele decreased with age within a single cage (e.g., because of drift and random chance) for non-significant SNPs. Specifically, we sampled 1000 frequency matched non-significant SNPs from the total list of non-significant SNPs for which we could assign ancestral/derived state, and then calculated the proportion of loci for which the derived allele decreased over time in one randomly drawn high sugar cage and one randomly drawn control cage, respectively. We performed this sampling 1000 times to get the null distributions shown in Figure 5F.

S10. Exploring the evolutionary history of lifespan-associated alleles

*Estimating allele frequency in Drosophila Nexus data*

To compare the allele frequency distribution of the North American population used in this study to that of African populations, we focused on all lifespan-associated (n=1007) and non-significant SNPs (n=6058) for which we had assigned an ancestral state (Table S12). We downloaded the sequence data for DPGP2 and DPGP3 from the *Drosophila* Genome Nexus website (https://www.johnpool.net/genomes.html). To identify our SNPs of interest in the Nexus dataset, we converted the coordinate position of the SNPs from *D. melanogaster* ref6 to *D. melanogaster* ref5 using the *Drosophila* Sequence Coordinate Converter on Flybase (https://flybase.org/convert/coordinates), and then tallied up the count of each allele (A, C, T, G or N) at each site. We excluded all SNPs whose coordinate positions did not fall on chromosome 2, 3 or X in *D. melanogaster* ref5 space, and all non-diallelic sites. For DPGP2, we excluded nine French samples and retained only African samples. The number of individuals used to estimate allele frequencies for each dataset are as following: DPGP2: chr2=113, chr3=128, chrX=124 and DPGP3: chr2=197, chr3=197, chrX=196. The final dataset consists of 868 significant and 4842 non-significant sites for which allele frequencies were determined in both, DPGP2 and DPGP3 datasets (Table S12).

*Testing whether lifespan-associated derived alleles show signatures of selection*

Given that we observed a bias towards derived alleles being lifespan-reducing, we focused on the set of 7062 SNPs for which we were able to assign ancestral state (Table S12) to ask about the evolutionary history of lifespan-associated derived alleles. To test whether lifespan-associated derived alleles might have experienced selection pressures during the evolutionary history of the North American population used here, we compared DAF between our experimental population and *D. melanogaster* populations from Africa where the species originated. For this, we focused on the set of SNPs for which we were able to estimate DAF in both African datasets (DPGP2 and DPGP3) in addition to ancestral state (n = 5710, Table S12). We first estimated ΔDAF as the difference between DAF in our experimental population at T0 and DAF in the African datasets. Then, we asked whether ΔDAF was different between lifespan-associated derived alleles and frequency matched non-significant sites. For this we used 1000 samples of non-significant sites sampled to match the frequency distribution of the lifespan-associated allele set. This test was done separately for the different sets of lifespan-associated alleles: lifespan-increasing, lifespan-decreasing, shared, and GxE derived alleles, and the results are shown in Figure 5C-D. Given that DAF do not differ between DPGP2 and DPGP3 datasets, we only show the results for DPGP2.

S11. Comparison of statistical power between this study and Mostafavi et. al. (2017)

We used simulations to compare the power of our study, which used data for 5,413 flies sampled at the beginning and end of the adult lifespan, to that of Mostafavi and colleagues (*13*), which used data for 57,696 humans sampled in middle age. These simulations were similar to those described above (see section *Comparison of two statistical approaches to test for allele frequency changes with age*) but were adjusted to parallel those described by Mostafavi and colleagues. In particular, we simulated genotype data for replicate datasets of 1000 loci with the following sample sizes and age distributions: 1) a dataset representative of Mostafavi and colleagues, with n=57,696 individuals sampled from the age distribution of the Genetic Epidemiology Research on Adult Health and Aging (GERA) cohort (as presented in Fig. S2 and

Data S1 of their publication) and 2) a dataset representative of our fly experiment but scaled to a human lifespan to be comparable to #1, with n=2,104 individuals sampled at age 0 and n=3,309 individuals sampled at the maximum allowable age in our simulations, 95 years. These sample sizes reflect our $T_0$ and $T_N$ samples sizes, respectively. For each of the two age structures (representing Mostafavi/GERA and this study), we performed replicate simulations in which we varied the starting minor allele frequency between 0.05 and 0.4; for all simulations, the percent decrease of the minor allele between age 0 and 95 was set to 20%, and the minor allele decreased at a constant rate across the lifespan.

We simulated genotypes for a given age class by drawing from a binomial distribution parameterized by the sample size and the minor allele frequency. For the fly simulations, these age classes were age 0 or age 95, and for the human simulations these age classes were the 5-year age bins used by Mostafavi and colleagues. For the fly simulations, we translated our simulated genotypes into count data obtained from low coverage sequencing by first sampling the total read counts from a negative binomial distribution parameterized by our real sequencing data (see section *Comparison of two statistical approaches to test for allele frequency changes with age*). We then simulated the number of minor allele reads for each individual at a given locus by drawing from a binomial distribution parameterized by the number of total reads and the focal individual's genotype. Mostafavi and colleagues analyzed array genotype data rather than low coverage sequencing data, so for the human simulations we did not translate genotypes into count data and assumed that genotypes were measured without error by the arrays.

For the fly simulations, count data were analyzed using a beta-binomial model with a fixed effect of age class implemented in the R package *aod* (*45*). We extracted the p-value associated with the age effect and used this to assess power. For the human simulations, we ran a binomial regression for each locus with the number of minor alleles as the outcome variable (coded as 0/2, 1/2, or 2/2) and a 14-way factor for age class as a fixed effects predictor variable as in (*13*). We also included a binary variable of "batch" as a fixed effect, which was randomly simulated with respect to both age and genotype. We then used a likelihood ratio test (LRT) to compare models with and without the age class factor, and extracted the p-value associated with the LRT to assess power. We also implemented the same statistical approach (using a binomial regression and LRT) for the fly simulations, to understand what power would look like if we had array genotype data rather than low coverage sequencing data. For all simulations, power was determined as the proportion of simulated loci with a p-value $<10^{-6}$.

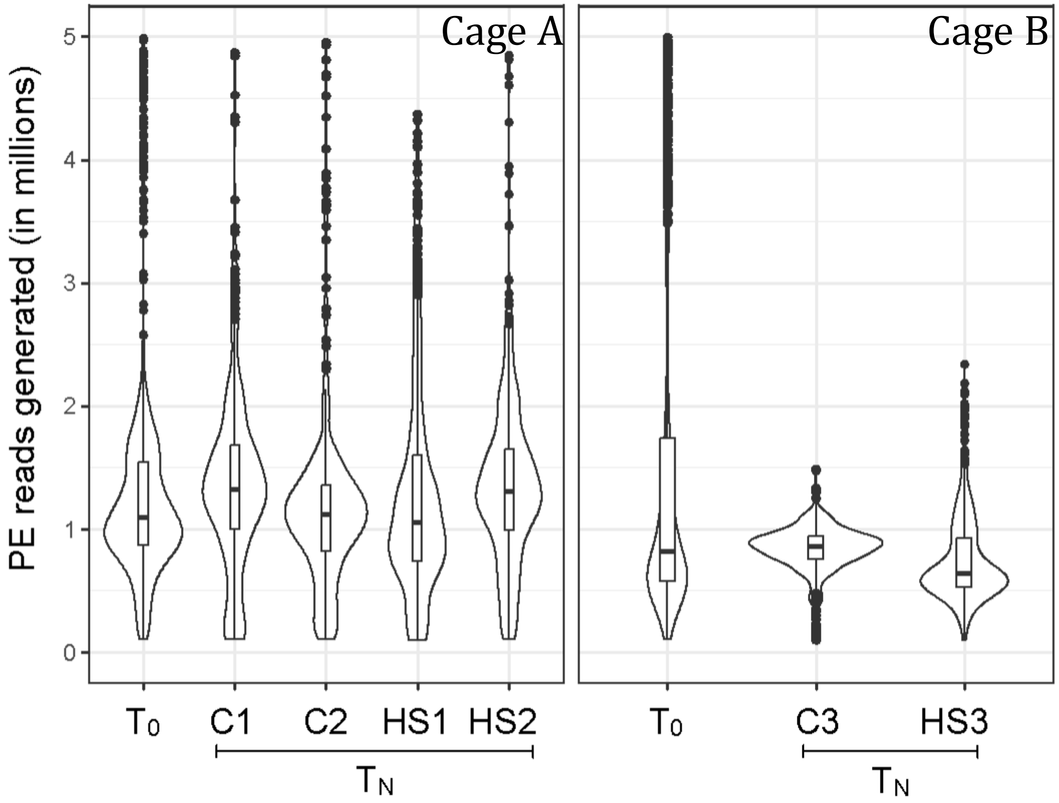


**Fig. S1. Mean read coverage for DNA libraries.** The y-axis represents the number of paired end (PE) reads generated per DNA library in millions. Samples are grouped by cage and time point.

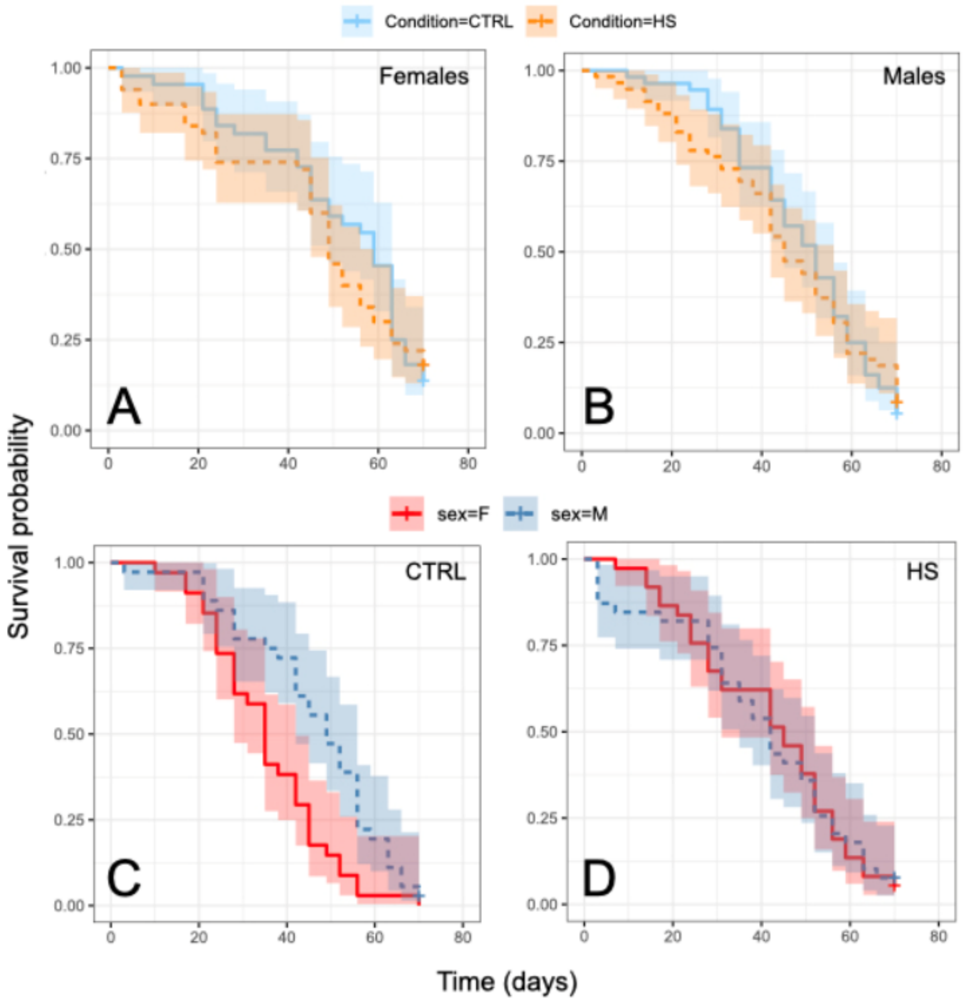


**Fig. S2. Sex and diet interact to affect survival.** Kaplan-Meier survival curves estimated for flies in vials of (A) only females or (B) only males on a CTRL or HS diet, respectively. When survival curves where estimated for males and females in mixed sex vials on (C) CTRL or (D) HS food, females exhibited significantly reduced longevity relative to males, but only on a CTRL diet. Full results from models testing for sex and diet effect on survival using a Cox proportional-hazards framework can be found in Table S5.

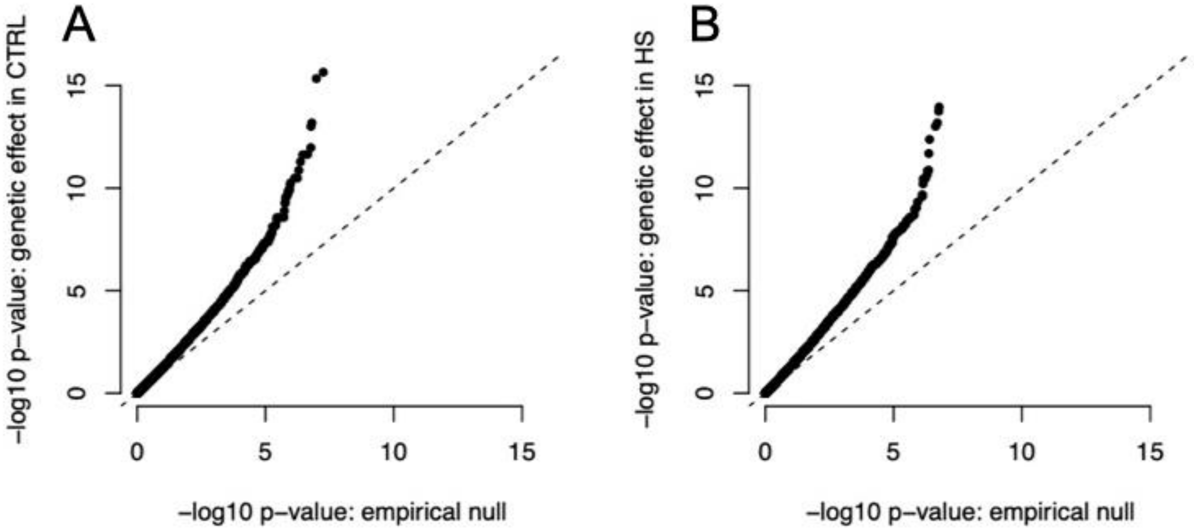


**Fig. S3. QQ-plots** comparing the empirical null distribution of p-values (obtained via permutations of the 3-way factor representing time and diet) to the observed distribution of p-values for the genetic effect on longevity on (A) a CTRL diet and (B) a HS diet.

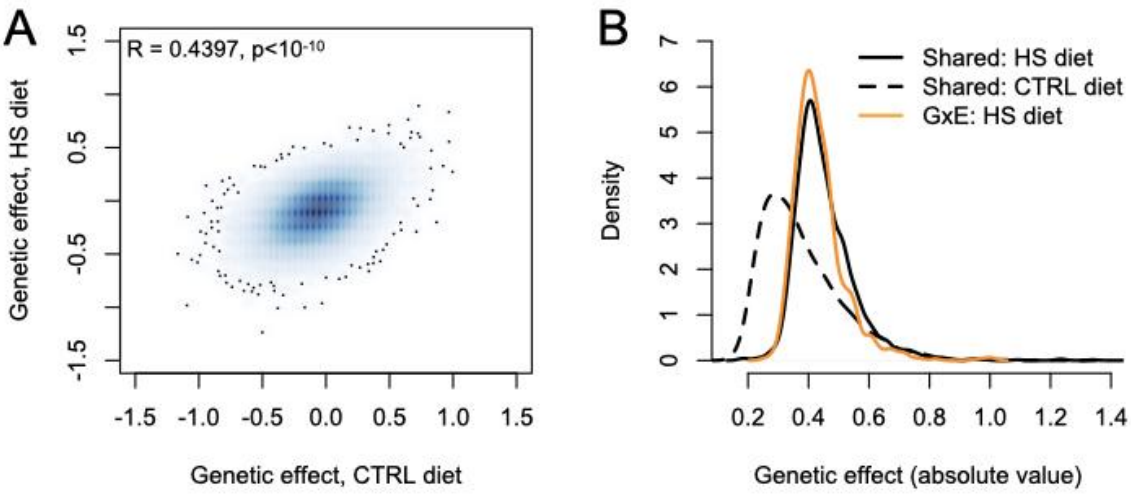


**Fig. S4. Comparison of the magnitude of the genetic effect on longevity across conditions.** **(A)** Across all tested sites, there is a strong correlation between the magnitude of the genetic effect on longevity estimated on a CTRL diet (x-axis) versus a HS diet (y-axis). **(B)** For sites with significant genetic effects on longevity that are shared between conditions (shared) or stronger in the HS environment (GxE), the magnitude of the longevity effect tends to be larger on a HS diet.

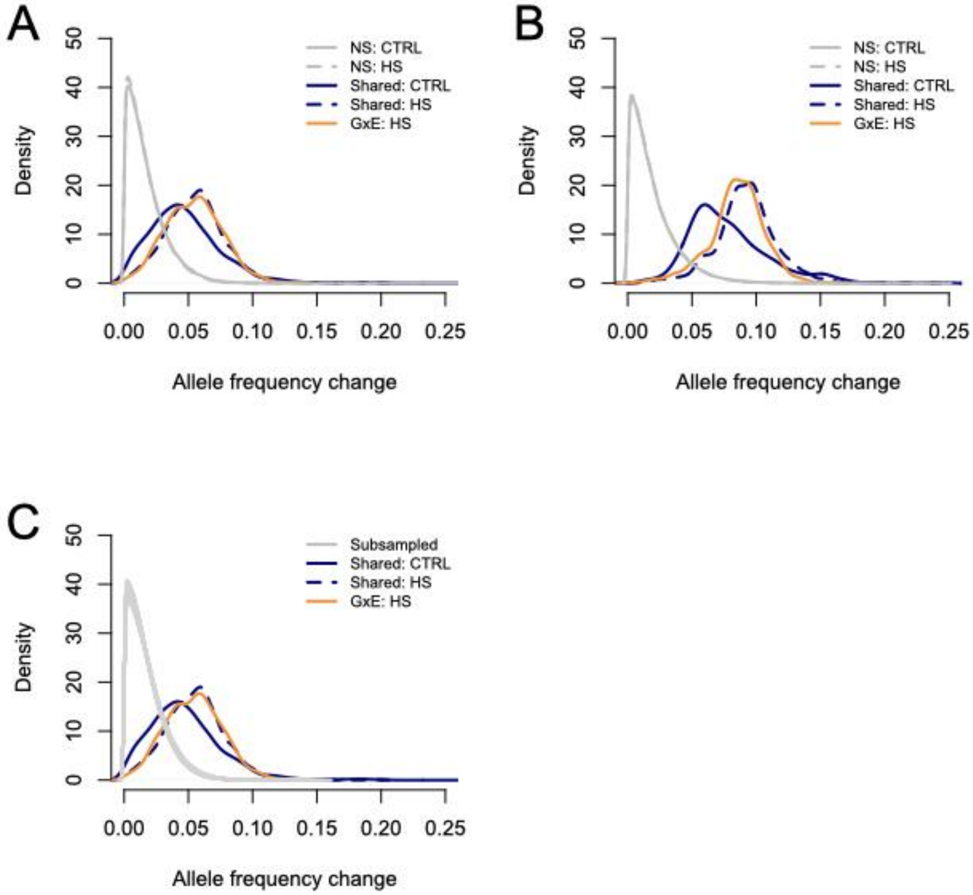


**Fig. S5. Allele frequency changes between $T_0$ and $T_N$ for SNPs with significant lifespan-effects**. **(A)** Absolute allele frequency change ($T_0$ vs $T_N$) on a CTRL or HS diet for sites with longevity effects shared between diets (dark blue solid lines) or HS-biased sites (orange line). The same distributions are also plotted for SNPs with no effects on longevity (grey). **(B)** Because there are other covariates in our experiment that also influence allele frequencies (i.e., sex, sequencing batch, and starting population cage), the "raw" observed allele frequency changes plotted in panel A may be downwardly biased or at the very least noisy. Therefore, we also translated our model fitted estimates of effect size (as shown in Figure 2C) into allele frequency changes. We used an approach similar to that of (*58*, *59*). Specifically, for each significant site, we calculated the allele frequency at $T_0$ by setting batch=1, sex=1, starting population pool=1, and CTRL and HS environment effects=0; we then summed the model fitted estimates of effect size multiplied by these values and passed the sum through a logistic function to obtain an allele frequency. To calculate the allele frequency at $T_N$ in the HS and CTRL environments, we performed parallel calculations where the CTRL or HS environment effects were set to 1. The distributions from these analyses are plotted here, with colors as in panel A. **(C)** Colored lines are the same as in panel A, but here the null distribution (grey lines) is derived by subsampling two sets of 1000 individuals with replacement at $T_0$ and calculating the absolute difference in allele frequency across all tested sites. This null distribution is plotted for 100 subsamples.

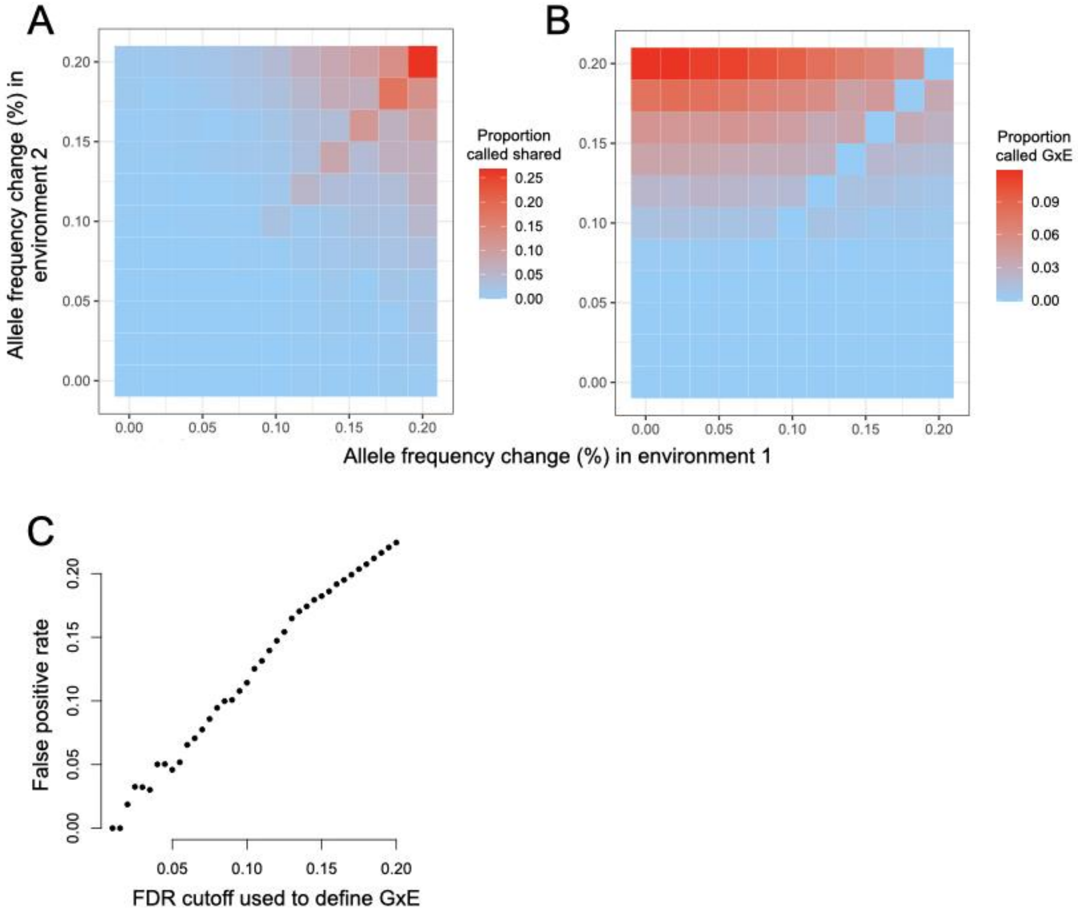


**Fig. S6. Simulations to understand our definitions of "shared" and "GxE" SNPs**. We classify SNPs as having "shared" effect sizes between environments (defined as FDR<10% in one environment and p<0.05 in the other) or "GxE" effects where the lifespan effect is stronger on one diet relative to the other ("GxE" defined as FDR<10% in one environment and p<0.05 in the other). To understand how this thresholding affected our results, we simulated 121 datasets of 1000 loci each where the lifespan reducing allele decreased by 0-20% with identical or varied effect sizes across two environments (sample sizes = 1000 individuals at $T_0$ and 1000 at each of the two $T_N$ end points). For each simulation, we recorded the number of loci that would be classified as **(A)** shared ($p<10^{-4.5}$ in one environment, similar to a 10% FDR in our real dataset, and p<0.05 in the other environment) or **(B)** GxE ($p<10^{-4.5}$ in environment #2 and $p>0.05$ in environment #1). See Text S6 for additional details. (**C**) The FDR cutoff used to define GxE effects (i.e., the threshold used to consider a site significant in one environment) versus the false positive rate estimated from permutations where the $T_N$ HS and $T_N$ CTRL labels were scrambled 10 times. See Text S6 for additional details.

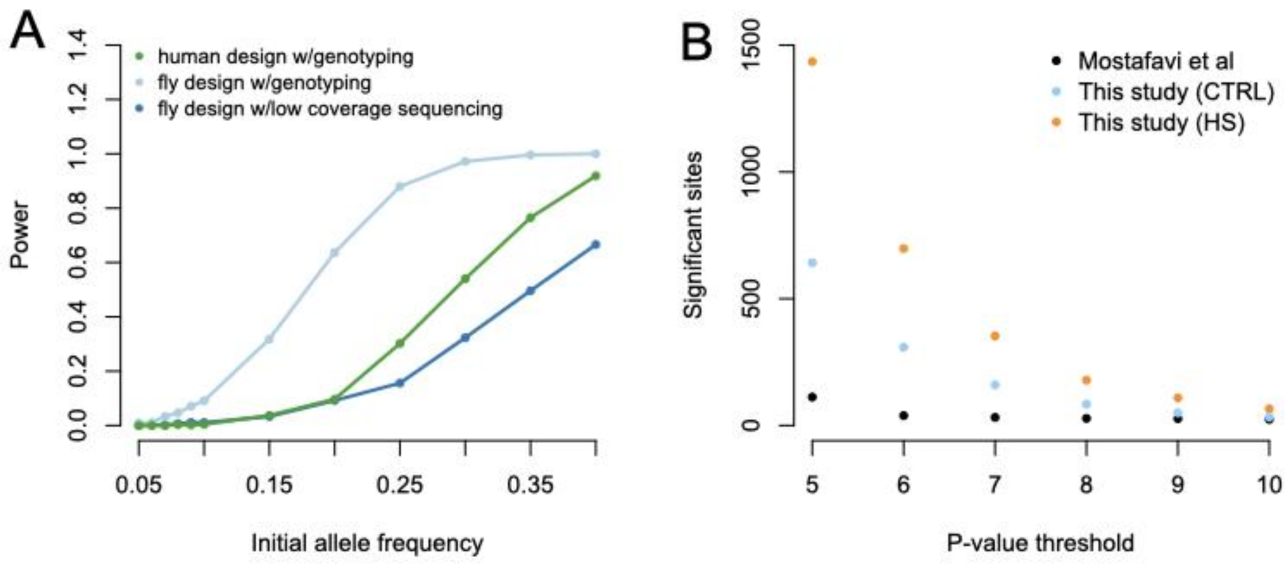


**Fig. S7. Power to detect a genetic effect on longevity using different study designs.** (A) Y-axis represents the proportion of simulated true positive sites (with a significant change in allele frequency across the lifespan) that are detected at $p<10^{-6}$. Shown are results from simulations that mimic the “fly” design used here, where individuals are sampled at the beginning and end of their life, versus the “human” design of Mostafavi *et al.* (*13*), where individuals were only genotyped in middle and old age. We performed simulations for the fly and human designs where true genotypes were estimated without error, as well as simulations where genotypes were derived from a low coverage (~1x) sequencing approach and therefore estimated with some error. The curves show simulation results for starting minor allele frequencies from 0.05-0.4, and effect sizes were always simulated as a 20% change in allele frequency from the youngest to the oldest age bin. Further details on the simulations and analysis of simulated data are provided in Text S11. Comparison of statistical power between this study and Mostafavi *et. al.* (*13*). (B) The number of longevity-associated SNPs found by this study versus Mostafavi *et al.* (*13*) at different nominal p-value thresholds.

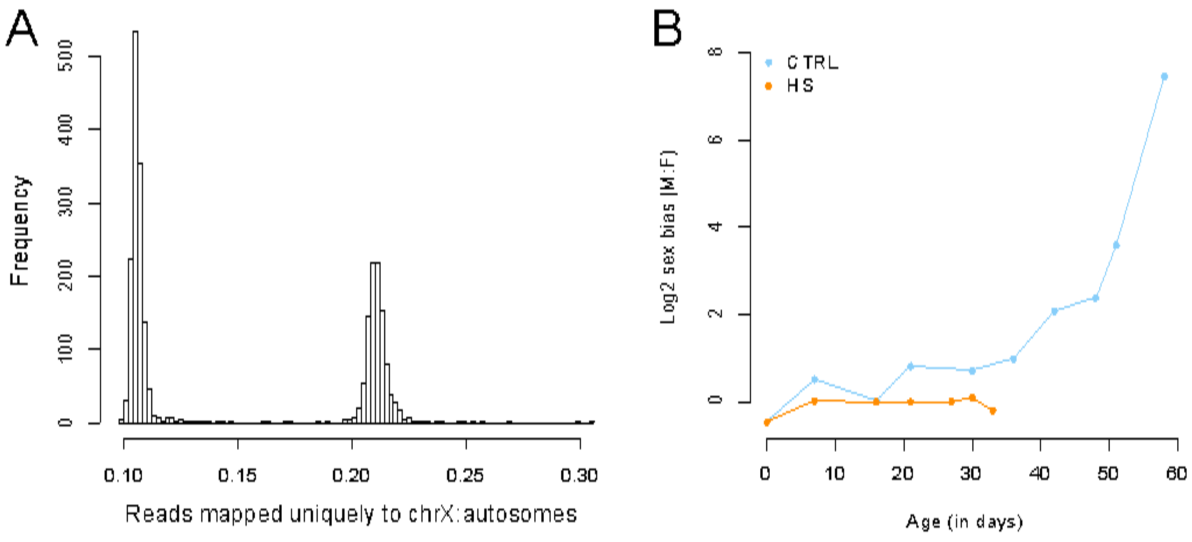


**Fig. S8. Sex determined from sequence data.** (A) Distribution of the proportion of reads that map uniquely to the X chromosome versus all of the autosomes. Samples for which this proportion was <0.13 were considered male, and samples for which this proportion was 0.18-0.25 were considered female. (B) Sex bias over time as estimated from the sequencing data for control and high sugar cages. To try to balance the representation of males and females in the sequencing efforts of the intermediate time points in CTRL cages, we included as many females as possible, and therefore the male bias in CTRL cages shown here is an underestimate. The sex bias estimated from sexing all flies collected at each intermediate time point (and not only the flies that were sequenced) is presented in Figure 1D.

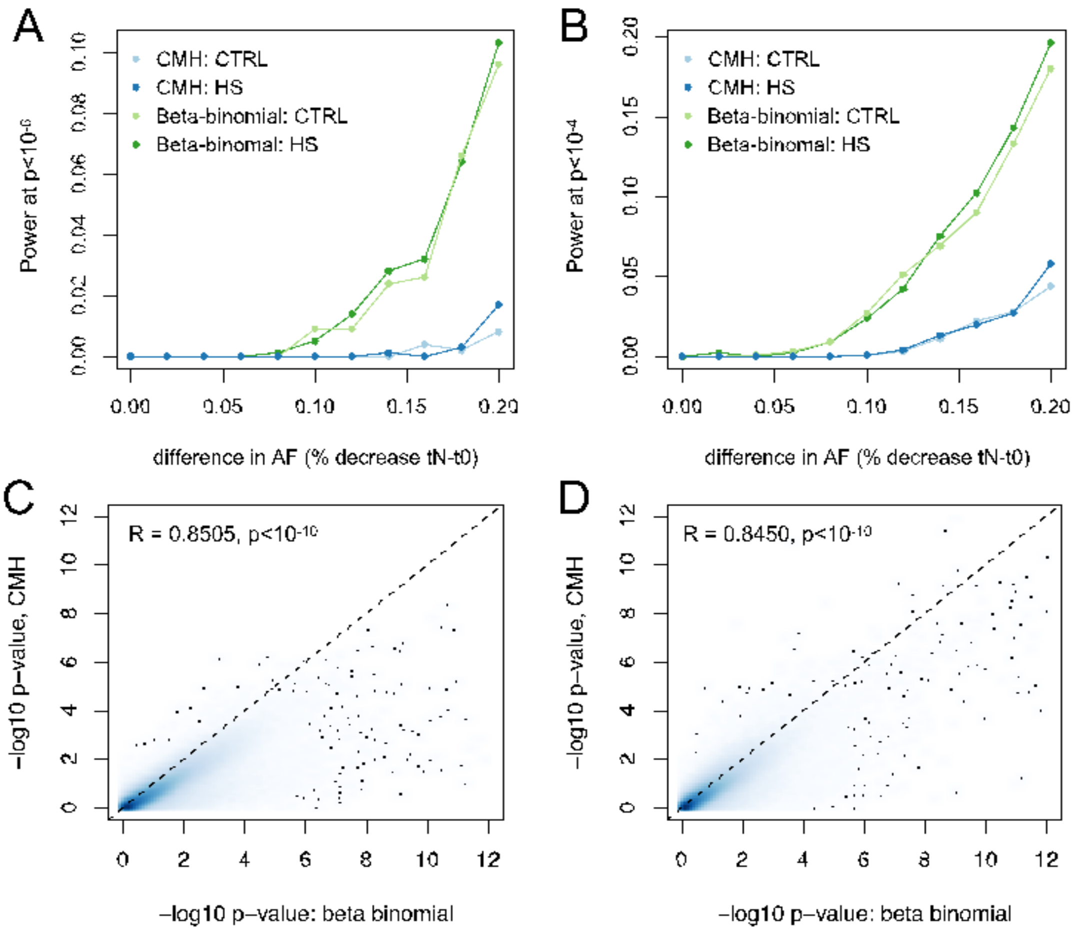


**Fig. S9. Power comparison between a Cochran-Mantel-Haenszel (CMH) and beta-binomial approach.** (A-B) The proportion of simulated true positives (i.e., SNPs with a significant effect on longevity) detected by a beta-binomial versus a CMH approach as a function of effect size. Different p-value thresholds are used to detect true positive SNPs in panels A and B as noted on the y-axis. Details on the simulations and analyses of simulated data are provided in the SI Methods (see Text S5. Comparison of two statistical approaches to test for allele frequency changes with age). Correlation between the -$\log_{10}$ p-value for a genetic effect on longevity from the real data, estimated in (C) CTRL or (C) HS conditions using a beta-binomial model (x-axis) or a CMH test (y-axis). As described in the main text, the p-values for the beta-binomial were estimated while controlling for covariates of sex, sequencing batch, and cage.
